# Genetic determinants of gene amplifications alter frequency and evolutionary trajectory of antibiotic resistance in *Staphylococcus aureus*

**DOI:** 10.1101/2025.04.10.647645

**Authors:** Kalinga Pavan T. Silva, Anupama Khare

**Affiliations:** Laboratory of Molecular Biology, National Cancer Institute, National Institutes of Health, Bethesda, MD 20892, USA

## Abstract

Gene amplifications are thought to be common in bacterial populations, providing a rapid reversible mode of adaptation to diverse stresses, including the acquisition of antibiotic resistance. We previously showed that the opportunistic pathogen *Staphylococcus aureus* evolves resistance to the dual-targeting fluoroquinolone delafloxacin (DLX) that inhibits both the DNA gyrase and DNA topoisomerase IV via gene amplifications of an efflux pump encoding gene *sdrM*. However, the pathways that control the formation or selection of gene amplifications, and consequently adaptive trajectories, remain understudied, especially in gram-positive bacteria like *S. aureus*. Here, we show that specific DNA repair and chromosomal separation pathways alter the frequency of formation and selection of gene amplifications in *S. aureus*. Through a screen of 36 mutants deficient in various DNA processes, we found that while *sdrM* amplification was still the almost universal path to DLX resistance, other mutations that increased *sdrM* expression reduced the selection frequency of *sdrM* amplifications, demonstrating the critical role of *sdrM* in DLX resistance. We found that similar to other bacteria, the formation and loss of *sdrM* amplifications required a functional RecA recombinase, but multiple other mutants in pathways required for amplifications in other species still exhibited frequent *sdrM* amplifications, suggesting that *S. aureus* may have alternate routes of amplification formation. Finally, mutants in the tyrosine recombinase XerC that is involved in chromosomal separation were deficient for *sdrM* amplifications, indicating that XerC is a novel modulator of amplification formation, maintenance, or selection. Thus, our work sheds light on genetic factors that alter gene amplification-mediated evolutionary trajectories to antibiotic resistance in *S. aureus* and can potentially unlock mechanisms by which such evolution of resistance can be inhibited.

## INTRODUCTION

Many antibiotics used to treat bacterial infections are becoming increasingly ineffective as clinical and environmental bacteria acquire resistance to them via horizontal gene transfer or de novo mutation-mediated evolution^1,2^. Such antimicrobial resistance is a major threat to global public health^3,4^. *Staphylococcus aureus*, especially methicillin-resistant *S. aureus* (MRSA), commonly acquires resistance to a broad range of antibiotics, and is a major cause of antibiotic resistant infections^5,6^.

While many instances of mutation-driven evolution of antibiotic resistance involve sequence alterations, gene amplifications can also lead to antibiotic resistance via a variety of mechanisms^7,8^. Gene amplifications have been previously implicated in resistance to multiple antibiotics in *S. aureus*. Duplications of the gene encoding the efflux pump *norA* led to ciprofloxacin resistance^9^, amplifications of the SCC*mec* cassette caused oxacillin resistance^10^, and other amplifications resulted in resistance against macrolides^11^ and intermediate resistance against vancomycin^12^.

Delafloxacin (DLX) is a dual-targeting antibiotic capable of inhibiting the activities of the DNA gyrase and DNA topoisomerase IV in multiple bacterial species including *S. aureus*^8,13^. Such antibiotics that target two or more components in the cell should theoretically lead to a lower frequency of resistance evolution, as cells would need mutations in multiple targets for resistance^14,15^. In an earlier study, we identified two distinct mechanisms leading to high DLX resistance in *S. aureus*. The more rapid and prevalent resistance evolutionary trajectory was via gene amplifications of the gene encoding an efflux pump SdrM that increased DLX efflux from the cell, while the second adaptive path, that we observed in the absence of functional *sdrM*, was through canonical mutations in both the DLX targets DNA gyrase and DNA topoisomerase IV^16^.

Previous work has shown a large reduction in duplication and amplification frequency in a *recA* null mutant in multiple bacterial species upon exposure to selective pressures^17,18^. Apart from RecA, other proteins involved in DNA double-strand break (DSB) repair, and some involved in other DNA metabolic processes are critical for the formation of gene amplifications in *E. coli*^19^. Our previous study showed that in the absence of a functional SdrM efflux pump, which is the amplification target, *S. aureus* did not evolve amplifications, and thus DLX resistance arose at a lower frequency via mutations in both the canonical target proteins DNA gyrase and topoisomerase IV^16^. However, other factors that may alter the formation or selection of gene amplifications, and thus evolutionary trajectories of antibiotic resistance, have not been characterized in *S. aureus*.

Here, we investigate genes and pathways involved in the selection of *sdrM* gene amplifications in *S. aureus*. We found that RecA is required for the formation of amplifications in *S. aureus*. In the absence of functional RecA, *S. aureus* required two steps to evolve DLX resistance, where the initial mutations partially compensated for the DNA damage sensitivity in the *recA* mutant, and were followed by mutations in *sdrM* or the canonical DLX targets. RecA also impacted the maintenance of the *sdrM* amplifications, as amplifications were more stable in the absence of functional RecA. Further, through a targeted screen of mutants in various DNA repair and metabolism pathways, we observed that *sdrM* amplifications remained the pervasive mechanism of DLX resistance, even in the absence of pathways that are required for the formation of amplifications in *E. coli*. However, the evolution of point mutations that increased expression of *sdrM* selected against *sdrM* amplifications, indicating that albeit less frequent, specific sequence alterations that achieve increased *sdrM* expression can lead to a lower selective advantage of *sdrM* amplifications, and highlighting the importance of SdrM function in conferring DLX resistance. Finally, we found that the widely conserved tyrosine recombinase XerC, involved in chromosomal separation, is a novel effector of the formation or selection of *sdrM* amplifications.

## RESULTS

### Resistance evolution via gene amplifications requires the recombinase RecA

The formation of gene amplifications is thought to be a two-step process where an initial duplication event is mediated by either homologous or non-homologous recombination, and is followed by higher order amplifications via RecA*-*mediated homologous recombination^8^. RecA is thus thought to be essential for the formation of gene amplifications^20^. To test whether this is also valid for gene amplifications in *S. aureus* and determine how the absence of RecA alters evolutionary trajectories of DLX resistance, we evolved three independent populations of a *S. aureus* JE2 *recA*::Tn mutant from the Nebraska Transposon Mutant Library (NTML) library^21^ for DLX resistance. The *recA*::Tn mutant is ∼56-fold more sensitive to DLX compared to the WT (**Supp. Fig. 1**). Fluoroquinolones cause DNA DSBs^22^ and, through a TUNEL assay we confirmed that DLX, a fluoroquinolone that targets both the DNA gyrase and topoisomerase IV enzymes, also leads to DNA damage (**Supp. Fig. 2**). This DNA damage likely underlies the significant DLX hypersensitivity of the DNA repair deficient *recA*::Tn mutant. We passaged the *recA*::Tn populations in increasing concentrations of DLX, starting from an initial DLX concentration ∼0.5x the MIC of the *recA*::Tn mutant, and continuing until a DLX concentration of at least 8 µg/mL. We performed whole genome sequencing (WGS) on populations from select intermediate passages as well as the terminal passage.

None of the three *recA*::Tn populations showed *sdrM* gene amplifications in any of the sequenced passages (**Fig. 1A**). Instead, mutations in the canonical targets DNA gyrase (*gyrA*, *gyrB*), and topoisomerase IV (*parC*, *parE*), and in and upstream of *sdrM* were prevalent in the later passages (**Fig. 1A** and **Supp. Table 1**). Interestingly, these mutations were selected during the evolution only once the evolving populations had reached a DLX resistance similar to that of the wild type (WT), while mutations in genes associated with DNA repair were observed in the early passages. Individual isolates from each population (R1, R2, and R3 from populations 1, 2, and 3 respectively) from a passage prior to the emergence of the canonical mutations had DLX resistance similar to that of the WT (**Fig. 1B).** Further, complementing RecA back into these isolates resulted in only an up to ∼2-fold increase in DLX resistance, compared to the ∼38-fold increase seen in the *recA*::Tn mutant (**Fig. 1C**), suggesting that these isolates may have recouped the intrinsic DLX resistance lost due to the nonfunctional RecA via mutations in genes different from the canonical targets or *sdrM*.

**Figure 1.**
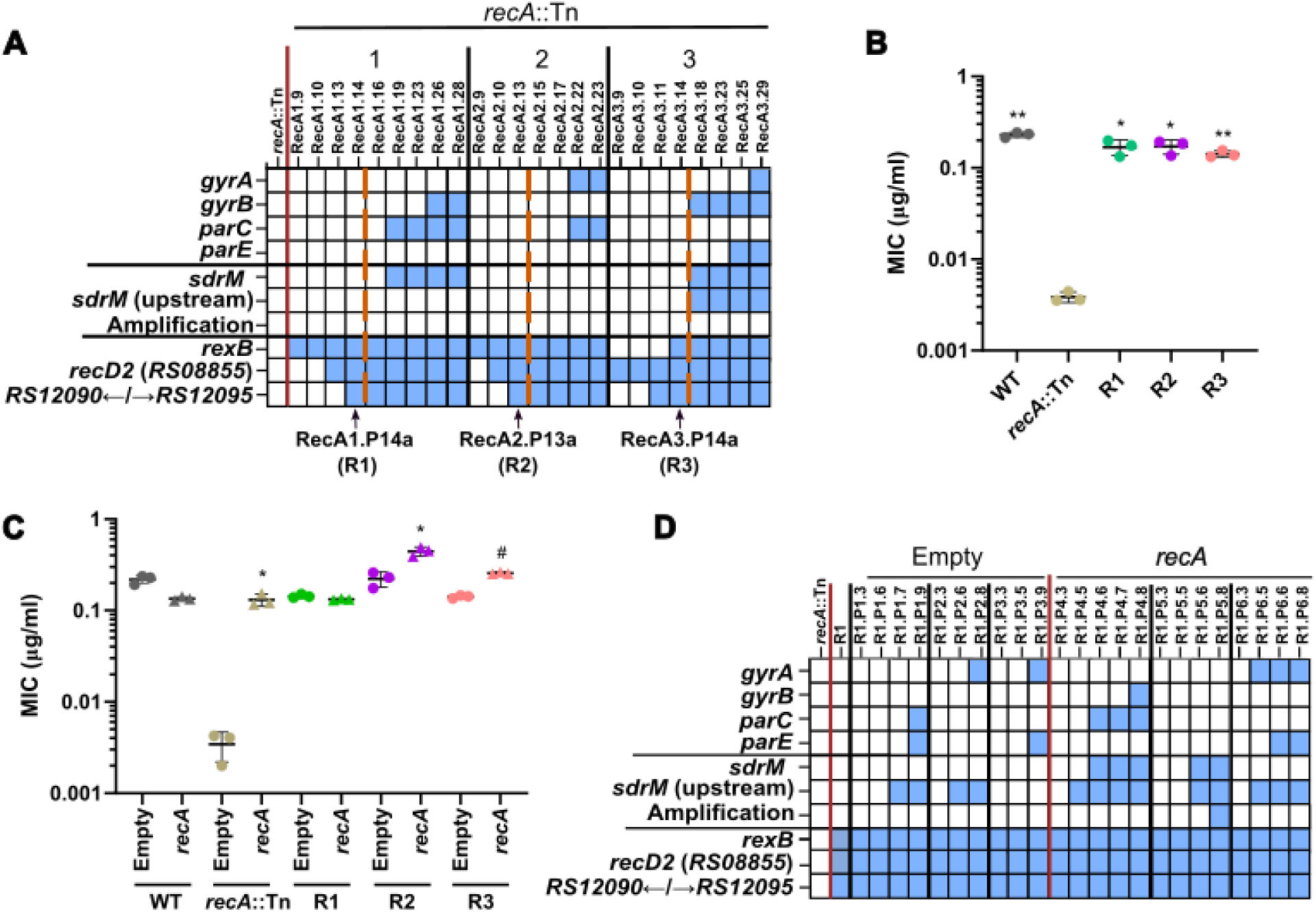
RecA is required for the formation of *sdrM* amplifications. **(A)** Common mutations and *sdrM* amplifications seen in three independently evolved populations of the *recA*::Tn mutant. Within each population, earlier to later passages are shown from left to right. Blue squares show the presence of an *sdrM* amplification or a mutation in the indicated genes. Dashed red lines indicate when the populations reached a resistance level similar to the WT. The populations from which the ‘intermediate isolates’ were selected are shown by black arrows. **(B)** DLX MICs of WT, the *recA*::Tn mutant, and the three intermediate isolates. **(C)** DLX MICs of WT, the *recA*::Tn mutant, and the three intermediate isolates, each carrying either an empty pKK30 plasmid, or one with the *recA* gene. **(D)** Presence of mutations (as shown by the blue squares) in the DNA gyrase and topoisomerase IV targets, *sdrM*, and DNA repair associated genes, as well as the *sdrM* amplifications, in three independently evolved populations each of the R1 intermediate isolate with either the pKK30 empty vector or one carrying *recA*. For each population, passages are shown chronologically from left to right. (**B, C)** Data shown are the mean ± standard deviation of three biological replicates. Significance is shown for comparison to **(B)** the *recA*::Tn mutant and **(C)** the respective empty vector strain as tested by Brown-Forsythe and Welch ANOVA tests followed by the Dunnett’s T3 multiple comparison’s test (* *p* < 0.05, ** *p* < 0.01, # *p <* 0.0001).

To further clarify if RecA is required for formation of the amplifications, we evolved three populations of the intermediate isolate R1 complemented with RecA, in parallel to three populations of R1 with an empty vector control. Given that the DLX MIC for R1 was similar to the WT, and it did not have any of the canonical mutations, we reasoned that if RecA was necessary for generating amplifications, re-introducing a functional RecA and selecting for DLX resistance should lead to *sdrM* amplifications. One of the three RecA-complemented populations showed amplifications, while no amplifications were seen in the empty vector control populations (**Fig. 1D and Supp. Table 2**), further validating that RecA is required for the emergence of *sdrM* amplifications under selection for DLX resistance.

### Absence of a functional RecA results in a two-step evolution of DLX resistance

RecA plays two important roles in DNA repair, first by activating the SOS response induced by DNA damaging agents, and second by acting as a critical effector of homologous recombination-mediated DSB repair^23,24^. We found that the intermediate isolates not only showed increased DLX resistance, but also had enhanced resistance against three other DNA damaging agents, a different fluoroquinolone ciprofloxacin (CPX)^25,26^, the anti-cancer drug doxorubicin (DOXO)^27^ and the chemotherapeutic mitomycin C (MMC)^28^, compared to the hyper-sensitive *recA*::Tn mutant (**Fig. 2A**).

**Figure 2.**
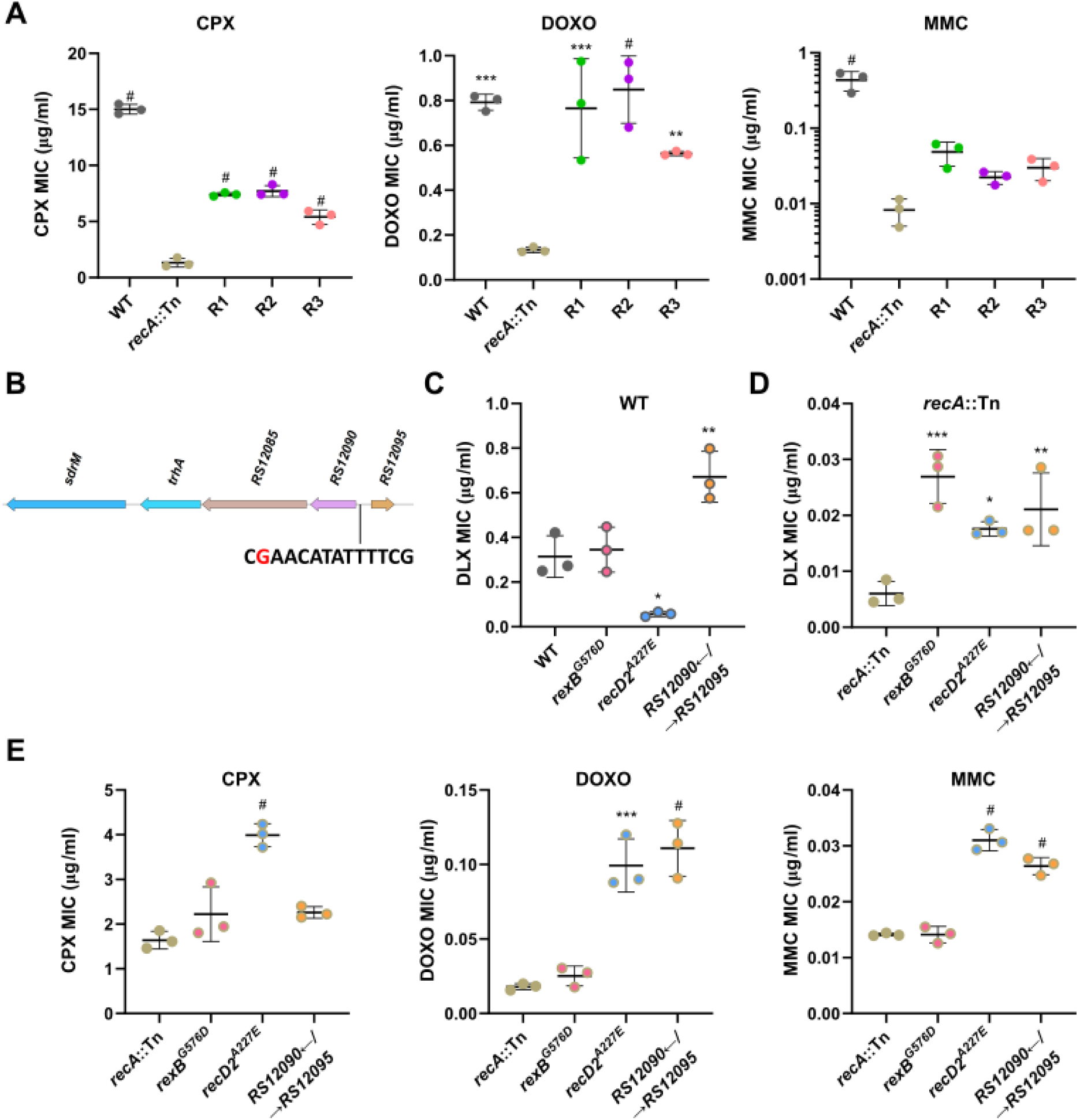
Mutations in *recA*::Tn intermediate isolates lead to DLX and DNA damage resistance. **(A)** MICs for ciprofloxacin (CPX), doxorubicin (DOXO), and mitomycin C (MMC) of WT, the *recA*::Tn mutant, and the three intermediate isolates. **(B)** The position of a G to A mutation (shown in red) in the putative LexA-binding site (shown in black) located between the genes *B7H15_RS12090* and *B7H15_RS12095*. **(C)** DLX MICs for WT and the indicated allele-replacement strains in the WT background. **(D)** DLX MICs for the *recA*::Tn mutant and the indicated allele-replacement strains in the *recA*::Tn background. **(E)** MICs for ciprofloxacin (CPX), doxorubicin (DOXO), and mitomycin C (MMC) of the *recA*::Tn mutant and the indicated allele-replacement strains in the *recA*::Tn background. **(A,C-E)** Data shown are the mean ± standard deviation of three biological replicates. Significance is shown for comparison to **(A)** the *recA*::Tn mutant, **(C-E)** the respective parental strain as tested by a one-way ANOVA with Dunnett’s test for multiple comparisons (* *p* < 0.05, ** *p* < 0.01, *** *p* < 0.001, # *p <* 0.0001).

Given that the intermediate isolates showed increased resistance to DLX and DNA damaging agents, but do not have canonical DLX resistance mutations, we investigated common mutations seen in these isolates, which were also seen in the early passages of the *recA*::Tn evolution (**Fig 1A**) and likely recoup some RecA functionality in dealing with DNA damage. All three isolates had coding sequence mutations in the genes encoding the helicase RexB, and a RecD2-like helicase, as well as an intergenic mutation upstream of the genes *B7H15_RS12090* and *B7H15_RS12095* (**Table 1**). RexB is involved in the processing of double stranded breaks (DSBs) during repair, similar to the activity of RecBCD in most gram-negative bacteria^23^. The two mutations observed in RexB, E48K and G576D, are both in residues that are conserved across species (**Supp. Fig. 3A**), and the E48 residue in the *Bacillus subtilis* homolog AddB has been implicated in binding to DNA^29^, indicating that mutations in these residues likely alter RexB function. While the role of the RecD2 helicase has not been characterized in *S. aureus*, RecD2 is present in other bacteria that lack RecBC and is thought to be involved in varied DNA repair processes^30–32^. In the intermediate isolates, we observed two different mutations, G360A and A227E, but the evolving populations showed numerous other coding sequence mutations, including A346D, T367P, T368T, T420M, D449G, Q480H, and N629K, which are all located in conserved residues (**Supp. Table 1 and Supp. Fig. 3B**). Given the diversity of residues, the effect of these mutations on DNA repair is unclear.

**Table 1:** Common mutations in the three *recA*::Tn evolved intermediate isolates.

| Strain | *rexB* | *recD2* | *RS12090*←\|→*RS12095*<br>at 2,306,688 |
| --- | --- | --- | --- |
| **R1** | E48K | G360A | G→A |
| **R2** | G576D | A227E | G→A |
| **R3** | G576D | A227E | G→A |

The third mutation we observed in all three intermediate isolates was an identical G to A substitution in the intergenic region between the adjacent divergently encoded genes, *B7H15_RS12090* and *B7H15_RS12095*. *B7H15_RS12090* is in the SOS regulon of *S. aureus* along with the two immediately downstream genes located in the same operon (*B7H15_RS12085* and *trhA*)^23^. The intergenic mutation lies within the predicted LexA binding site in this region^25^ (**Fig. 2B**) and may thus lead to the upregulation of these three genes.

To test the role that each mutation might be playing in conferring increased DLX resistance, we transferred *rexB^G576D^* (*rexB\**), and *recD2 ^A227E^* (*recD2*\*), which were found in two intermediate isolates each, as well as the intergenic mutation between *B7H15_RS12090* and *B7H15_RS12095* individually to both the WT and *recA*::Tn backgrounds. We observed that while only the intergenic mutation conferred a ∼2-fold increase in DLX resistance in the WT background, (**Fig. 2C**), all three mutations individually led to elevated DLX resistance in the DLX-hypersensitive *recA*::Tn mutant background (**Fig. 2D**), accounting for their selection during the evolution. Further, in the *recA*::Tn background, while the *recD2\** allele-replacement strain was resistant to all three DNA damaging agents, and the intergenic mutation led to resistance against DOXO and MMC, *rexB*\* did not cause significant resistance against any of the agents, indicating a specific effect on DLX resistance in the absence of functional RecA (**Fig. 2E**). Thus, in the absence of *sdrM* amplifications, the *recA*::Tn mutant evolves DLX resistance via a two-step process: initial DNA repair mutations that partially compensate for its DNA damage sensitivity, followed by point mutations in the DNA gyrase, DNA topoisomerase IV, and *sdrM*.

### RecA is required for loss of the *sdrM* amplifications

To test whether RecA plays a role in the instability of *sdrM* amplifications, we introduced the *recA*::Tn mutation to an evolved strain from our previous study, 1.7a, which had *sdrM* amplifications leading to a ∼140-fold increase in DLX resistance compared to the WT^16^. The 1.7a *recA*::Tn strain had a ∼17-fold reduction in DLX resistance compared to the parental 1.7a strain, and a ∼266-fold increase compared to the *recA*::Tn strain (**Fig. 3A**). We passaged this strain in different concentrations of DLX, and found that unlike what we had previously seen in the 1.7a mutant^16^ (shown here in the left panel of **Figure 3B**), the copy number of *sdrM* did not show significant variability in the 1.7a *recA*::Tn mutant (**Fig. 3B**), suggesting that RecA likely plays a role in the expansion of *sdrM* amplifications. Further, we passaged 1.7a and 1.7a *recA*::Tn in DLX-free media for five passages and found that the normalized *sdrM* copy number compared to the initial passage remained significantly higher in the 1.7a *recA*::Tn strain compared to in 1.7a, where the copy number reached ∼1 indicating an almost complete loss of amplifications (**Fig. 3C**).These data validate that RecA is critical for the loss of *sdrM* amplifications. Given the stability of the *sdrM* amplifications in the 1.7a *recA*::Tn strain, we tested its growth in DLX-free media to determine the effect of the *sdrM* amplifications on fitness. We found that while 1.7a showed similar growth compared to the WT, and the *recA*::Tn mutant grew slower than those strains, the 1.7a *recA*::Tn mutant showed substantially reduced growth even compared to the *recA*::Tn mutant, indicating a significant fitness cost of the *sdrM* amplifications, at least in the absence of functional RecA (**Fig. 3D and Supp. Fig. 4**).

**Figure 3.**
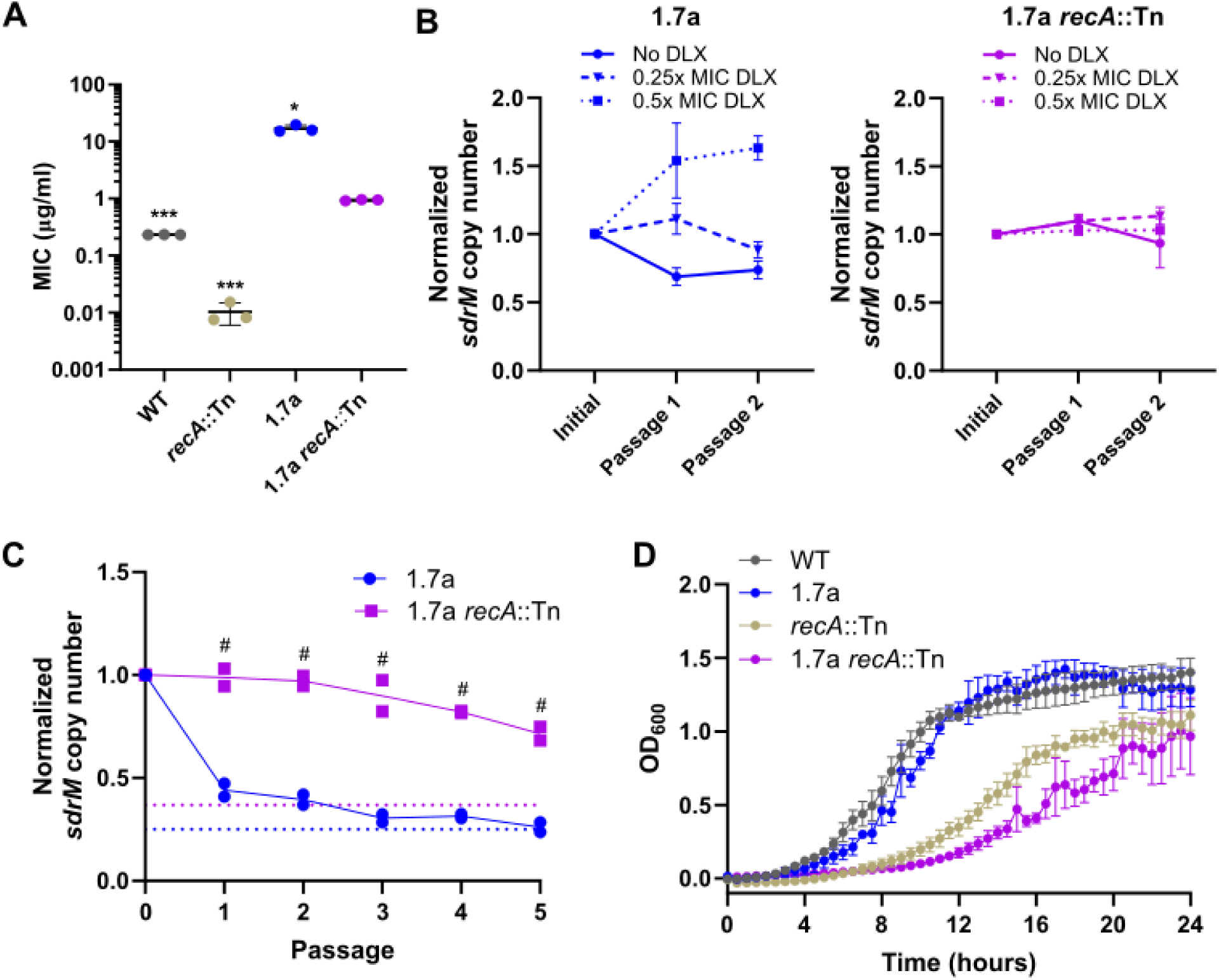
Amplifications are stabilized in the absence of RecA and have a fitness cost. **(A)** DLX MICs for WT, and 1.7a, as well as *recA*::Tn mutants in both backgrounds. Data shown are the mean ± standard deviation of three biological replicates. **(B)** Normalized *sdrM* copy number (normalized to the copy number in the initial culture) for 1.7a (left) and 1.7a *recA*::Tn (right) upon passaging in either no DLX, or in two different DLX concentrations, corresponding to ∼0.25x and ∼0.5x the respective MICs. The data for 1.7a are from our previous study^16^ licensed under the <u>Creative Commons Attribution 4.0 International License</u> and are shown here as *sdrM* copy number normalized to the initial value (instead of absolute *sdrM* copy number as in the original figure). Data shown are the mean ± standard deviation of two independent passaging experiments. **(C)** Normalized *sdrM* copy number (normalized to the copy number in the respective initial culture) for 1.7a and 1.7a *recA*::Tn upon extended passaging without DLX. The dotted lines denote an absolute copy number of 1 for the respective samples. Data from two independent passaging experiments are shown. **(D)** Growth curves shown as OD_600_ measurements of WT, and 1.7a, as well as *recA*::Tn mutants in both backgrounds. Data shown are the mean ± standard error of the mean of three biological replicates. Significance is shown for comparison to **(A)** the 1.7a *recA*::Tn strain, and **(C)** the respective 1.7a passage as tested by **(A)** Brown-Forsythe and Welch ANOVA tests followed by the Dunnett’s T3 multiple comparison’s test or **(C)** a two-way ANOVA with Sidak’s multiple comparisons test (* *p* < 0.05, *** *p* < 0.001, # *p <* 0.0001).

### Mutant library screen shows that evolution of DLX resistance leads to prevalent *sdrM* amplifications with diverse junctions

The absence of functional SdrM alters DLX resistance trajectories by requiring dual target mutations in the DNA gyrase and topoisomerase IV^16^, while the absence of RecA leads to a two-step evolution of DLX resistance (**Figs. 1 and 2**). To identify other genes that modulate the formation or selection of *sdrM* amplifications, and thereby the evolutionary trajectory of DLX resistance, we considered genes involved in DNA repair and recombination, the SOS regulon, and chromosomal separation, and those located in the genomic neighborhood of *sdrM*. We identified 33 genes of interest that had available loss of function transposon mutants in the NTML^21^ (**Supp. Table 3**). We also constructed two mutants that had *lexA* alleles (*lexA^S130A^* and *lexA^G94E^*) previously reported to encode uninducible versions of *lexA*, thereby inhibiting the SOS response^25,33,34^. We determined the DLX MICs of the corresponding mutants (**Supp. Fig. 1**) and for most mutants evolved three independent populations in increasing DLX concentrations, starting from ∼0.5x the respective MIC. We initially evolved the populations up to a DLX concentration of 2 µg/ml and tested for the presence of amplifications using WGS. For the populations without amplifications, evolution was continued until a DLX concentration between 8-16 µg/ml.

For the two mutants with the lowest MICs, *rexA*::Tn and the previously described *recA*::Tn, we realized that the initial mutations were in genes related to DNA repair, and canonical DLX resistance mutations arose later, once the evolving populations had an MIC similar to the WT (**Fig. 1A and Supp. Fig. 5**). We reasoned that for other mutants with similarly low MICs, the first mutations selected would likely recoup their DNA repair defect, and only subsequently select for canonical target or *sdrM* mutations, or *sdrM* amplifications. Therefore, given our goal of identifying genes required for amplification formation, for other mutants with very low MICs (*xerC*::Tn and *recG*::Tn), we evolved one population until its MIC was similar to the WT and then split that intermediate population into three independent populations for further evolution.

Given the large numbers of populations evolved, we first analyzed the general trends for DLX resistance in *S. aureus*. Of the 105 independent populations evolved, 89 had *sdrM* amplifications, reinforcing the finding that *sdrM* amplifications comprise a major evolutionary path to DLX resistance. 103 distinct genomic segments around *sdrM* were amplified, each found in only one evolved population, with the exception of two amplified segments that were found in two different independently evolved populations each (**Supp. Table 4**). The amplified segments were diverse both in their lengths and exact genomic coordinates of their ends (**Fig. 4A-4C**). While most of the upstream ends of the fragments were located in a ∼4.7kb kb region starting ∼50bp upstream of *sdrM*, all the downstream ends were located in a ∼6 kb region starting ∼3.2 kb away from *sdrM* (**Fig. 4D**). These ends were located almost exclusively in a tRNA-rRNA cluster, located four genes downstream of *sdrM*, resulting in these four genes also being present in most amplifications. Our previous work had showed that within these amplifications, DLX resistance depends solely on the overexpression of *sdrM*^16^. The prevalence of amplification ends in the tRNA-rRNA cluster, skipping the intermediate genes, thus indicates that the tRNA-rRNA cluster may be a hot-spot for the formation of amplifications. Finally, the ends of the amplifications had limited homology (**Fig. 4E**), indicating a role of non-homologous recombination in the formation of these junctions, as observed in the WT in our previous study^16^.

**Figure 4.**
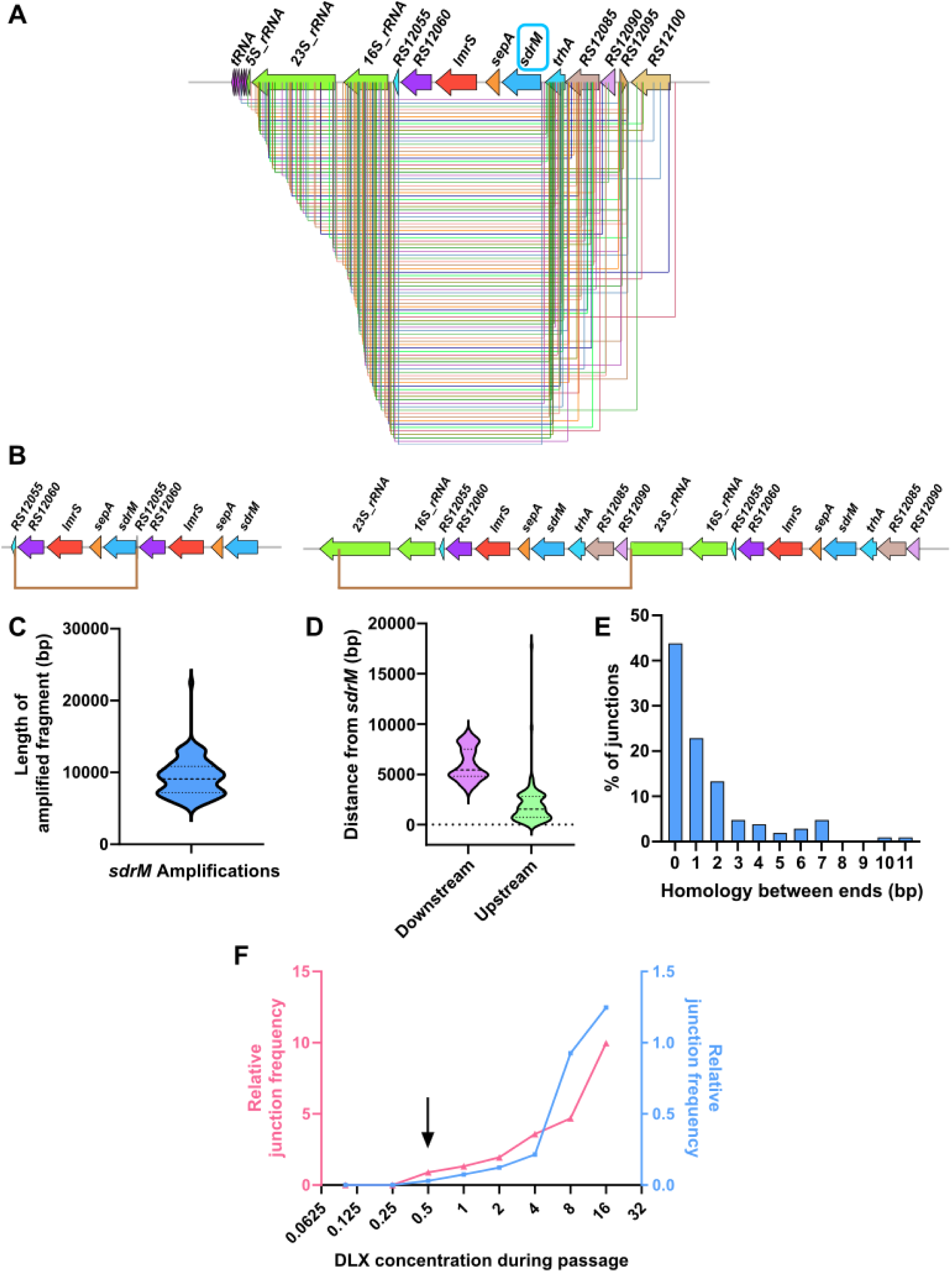
Amplifications of *sdrM* are highly prevalent, diverse, non-homologous, and originate in an rRNA-tRNA locus. **(A)** All distinct amplified fragments containing *sdrM* seen in our evolved populations. The terminus downstream of *sdrM* in almost all the fragments is in the tRNA-rRNA locus. **(B)** Two examples of *sdrM* amplifications each showing two copies of the amplified fragment (bordered by the brown lines). The one on the left is among the shortest amplified fragments, whereas the one on the right is significantly longer. **(C)** Length distribution of the amplified fragments containing *sdrM*. **(D)** Distribution of the distances of the downstream and upstream ends of the amplified fragments from the coding sequence of *sdrM*. The downstream ends are significantly further away as they are concentrated in the rRNA-tRNA locus, while the upstream ends start immediately upstream of *sdrM*. **(E)** Distribution of the length of homology between the two ends of the amplified fragments indicates a lack of significant homology. **(F)** Relative frequency of novel junctions representing a duplication (compared to *rpoC* levels), as measured by qPCR, in genomic DNA from successive passages of two independently evolved WT populations. For both populations, the junctions were detected in the passage containing 0.5 µg/mL DLX (shown by the black arrow).

Given the numerous distinct amplifications we found, we wanted to determine whether these could be detected in the unselected parental strains. We first measured the detection limit for two novel junctions that were selected for in one WT population each to be 1 in ∼10^4^-10^5^ cells (**Supp. Fig. 6**). Further, we found that in these two WT populations, the junctions were first detected in populations being passaged at 0.5 µg/ml DLX, which is ∼2x the WT MIC (**Fig. 4F**), but not in the previous passages. Thus, if these junctions, which represent at least a duplication, were present in the unselected parental strain, they must exist at a frequency lower than 1 in 10^4^-10^5^ cells. To identify any other junctions in this region in the unselected WT strain, we used an inverse PCR-like strategy, in conjunction with TOPO TA cloning (**Supp. Fig. 7A**). We identified three different junctions from three independently grown WT cultures (**Supp. Fig. 7B**), neither of which were present in any of our evolved populations. However, we could not detect them using junction-specific primers, indicating that their frequency in the population was less than 1 in ∼10^4^-10^5^, or they could be an artifact of the cloning.

### Mutations increasing *sdrM* expression are associated with a lack of *sdrM* amplifications

We next examined the mutations observed during the evolution, focusing on common mutations observed in more than one mutant background (**Supp. Table 5**). Apart from amplifications of *sdrM*, evolved populations from almost all mutants also showed increased copy number of a ∼43 kb genomic region (**Fig. 5A**). Using PHASTER, an online tool that identifies and annotates prophages^35,36^, we determined that this region encodes an intact prophage (**Supp. Fig. 8**), suggesting that exposure to increasing DLX concentrations led to induction of this prophage. Similar to our previous study^16^, many evolved populations contained mutations in the canonical DNA gyrase and topoisomerase IV targets and in *sdrM* (**Fig. 5A**). Apart from the previously identified coding sequence mutations in *sdrM* (A268S and Y363H) that increase DLX efflux and resistance^16^, we also observed additional *sdrM* mutations that emerged including T180I, Y363F, Y363N, K390I, K390N, K390T, and an in-frame 3 bp deletion resulting in removal of the threonine at position 8 (**Supp. Table 5**), which may be novel alleles of *sdrM* that confer DLX resistance. A few other common mutations were coding sequence and intergenic upstream alterations in genes associated with DNA damage and repair (*recD2*, *dinG*)^37,38^, bacterial transcription (*nusA)*^39^ and antibiotic resistance (*norA*)^9^, which may play a role in DLX resistance at least in the specific mutant backgrounds where they evolved.

**Figure 5.**
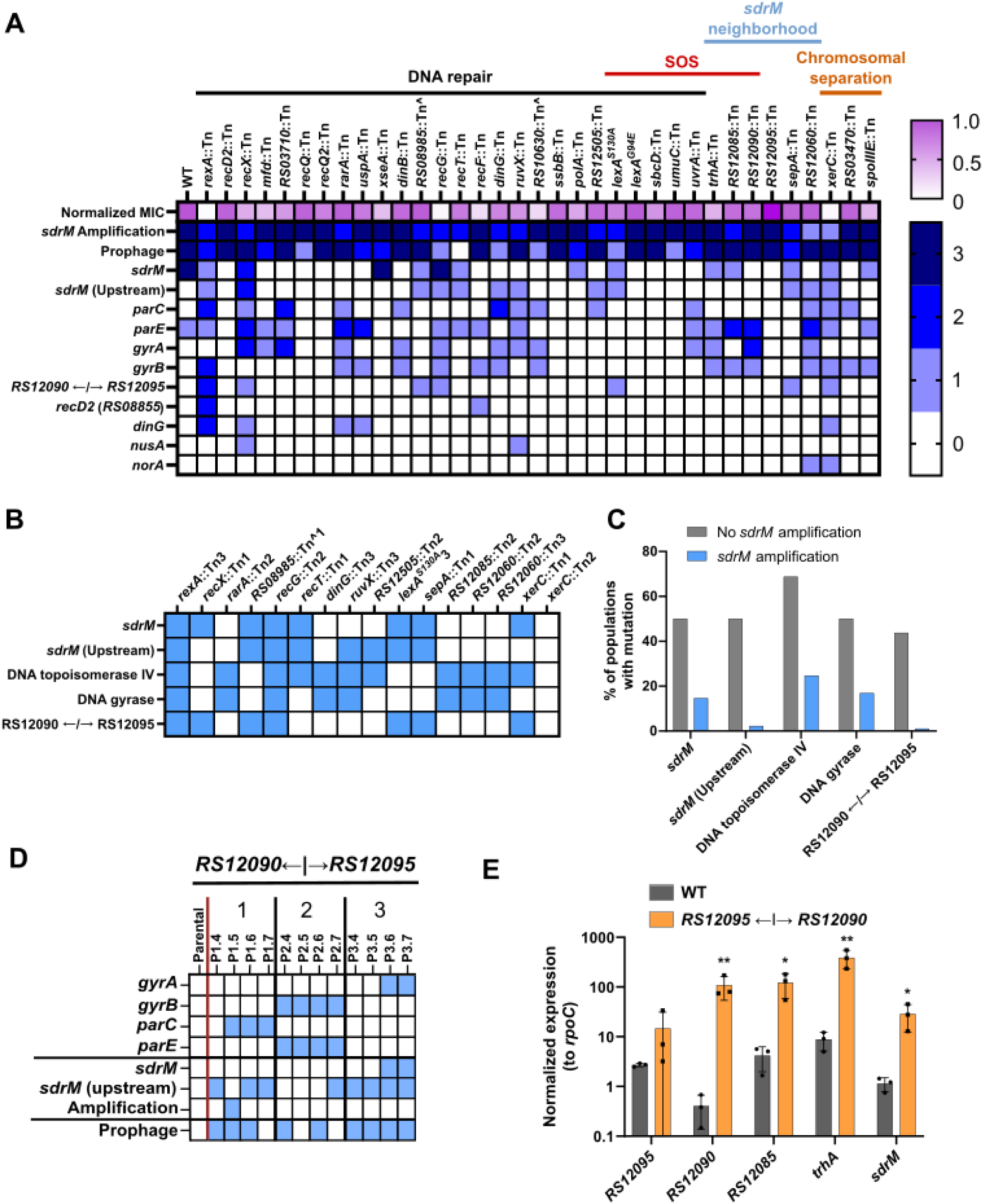
Amplifications of *sdrM* are prevalent upon selection of DLX resistance but negatively associated with mutations increasing *sdrM* expression. (A) Relative MICs (normalized to the WT) and the number of independently evolved populations that showed the mutations in the indicated genes, as well as the *sdrM* amplifications and increased prophage coverage, for WT and the screened mutants (a complete list of mutants is in **Supp. Table 3**). **(B)** Common mutations (indicated by the blue squares) seen in the independently evolved populations that did not contain *sdrM* amplifications. **(C)** Percentage of the populations with or without *sdrM* amplifications that had mutations in the indicated genes. **(D)** Common mutations seen in three independently evolved populations of an allele-replacement strain carrying the intergenic mutation between the genes *B7H15_RS12090* and *B7H15_RS12095*. Presence of mutations in the corresponding genes is indicated by a blue square. For each population, passages are shown from left to right in chronological order. **(E)** Normalized expression (compared to *rpoC*) of the indicated genes in the WT and the allele replacement strain carrying the intergenic mutation in the LexA-binding site. Data shown are the mean ± standard deviation of three biological replicates. Significance is shown for comparison to the respective WT sample as tested by ratio paired *t-*tests (* *p* < 0.05, ** *p* < 0.01).

In evolved populations without *sdrM* amplifications we observed mutations in the DLX targets (DNA gyrase and topoisomerase IV), coding and upstream mutations in *sdrM*, and mutations in the intergenic region upstream of the genes *B7H15_RS12090* and *B7H15_RS12095* (**Fig. 5B**). For the intergenic region, most strains had the same G to A substitution in the LexA-binding site seen in the *recA*::Tn populations (**Fig. 2B**), while one strain had a single base-pair deletion at the same position. While the frequency of all these mutations was higher in the populations without *sdrM* amplifications (**Fig. 5C**), mutations upstream of *sdrM*, and the intergenic mutation upstream of *B7H15_RS12090* and *B7H15_RS12095* were almost exclusively seen in evolved populations without *sdrM* amplifications. Further, in an allele-replacement strain carrying the intergenic mutation in the putative LexA-binding site, only one of three populations selected for DLX resistance transiently evolved an *sdrM* amplification (**Fig. 5D and Supp. Table 6**). Thus, the presence of the intergenic mutation upstream of *B7H15_RS12090* and *B7H15_RS12095* reduces the frequency of selection of *sdrM* amplifications upon DLX exposure.

Three genes in that region are thought to be in the SOS regulon, and their expression is likely to be affected by mutations in the LexA binding site^25^. We found that the expression of these genes as well as the adjacent genes *sdrM* and *B7H15_RS12095* is significantly higher in the presence of the LexA-binding site mutation, compared to the WT strain (**Fig. 5E**), indicating that *sdrM* expression (as well as that of *B7H15_RS12095*) may also be controlled by the SOS response.

Further, in a mutant containing an *sdrM* intergenic mutation at the-164 position along with the A268S coding sequence mutation that we had previously studied^16^, *sdrM* expression was significantly higher compared to a mutant containing only the A268S mutation (**Supp. Fig. 9**). The mutations we see in the intergenic region in our evolved populations are mostly clustered between positions-145 to-148 and-161 to-164 (**Supp. Table 5**) and accordingly may also confer DLX resistance by increasing *sdrM* expression. Thus, mutations that increase *sdrM* expression likely reduce the selective advantage of *sdrM* amplifications, providing alternate trajectories of DLX resistance evolution.

### XerC is a novel modulator of *sdrM* gene amplifications

Of the 35 mutants tested, 33 had amplifications in at least two of the three evolved populations (**Fig. 5A**), suggesting that the genes that are mutated in these strains may not play a major role in amplification formation. These include RexA, which together with RexB, is thought to play a role in DSB repair, which has been implicated in the formation of amplifications in *E. coli*^19^. In two of the *rexA*::Tn evolved populations, mutations associated with the RecD2 DNA helicase, DinG exonuclease, RecJ exonuclease, SbcCD endonuclease, or RNA polymerase subunit RpoC, were observed before selection of *sdrM* amplifications, or mutations in the canonical targets, *sdrM*, or LexA binding site (**Supp. Fig. 5**). This indicates that as seen in the *recA*::Tn populations, the *rexA*::Tn strains were recovering DNA repair function prior to specific DLX resistance. Unlike the *recA*::Tn evolutions, we did not observe mutations in RexB in these strains, likely because RexB is unable to participate in DNA repair functions in the absence of functional RexA.

We also tested mutants in multiple DNA polymerases, helicases, and nucleases that are thought to be involved in DNA repair and recombination^23,40^, and most of these mutants evolved amplifications in at least two of the three evolved populations. Further, the two uninducible LexA alleles did not alter the formation of amplifications, indicating that the SOS response does not play a significant role in the formation of gene amplifications, similar to what has been reported in *E. coli*^41^. However, two mutants, *xerC*::Tn, and *B7H15_RS12060*::Tn had only one independent population with *sdrM* amplifications, indicating a potential defect in amplification formation.

XerC is a universally conserved tyrosine recombinase that is critical for the separation of chromosomes^42^, and *B7H15_RS12060* encodes a putative P-Loop NTPase that is located in the neighborhood of *sdrM*. To further test whether these genes played a role in amplification formation, we conducted a secondary screen. We evolved 12 independent populations of each mutant in a 96-deep well plate for 14 days, where each population was passaged daily in a serial dilution of DLX, and the population growing in the highest DLX concentration was propagated to the next passage (**Supp. Fig. 10**). As controls, we also performed this evolution for the WT, and for a mutant (*B7H15_RS12085*::Tn) that showed *sdrM* amplifications in two out of three populations in our initial evolution (**Supp. Table 7**).

For both the WT and the control mutant, nine out of the 12 evolved populations contained *sdrM* gene amplifications (**Fig. 6A** and **Supp. Table 7**), indicating that mutants that had two out of three evolved populations with amplifications in the original tube evolution may not have defects in amplification formation. For the *B7H15_RS12060*::Tn strain, only five out of twelve populations evolved gene amplifications, while none of the *xerC*::Tn mutant populations had gene amplifications (**Fig. 6A**). Further, in this resistance evolution protocol, evolved populations without *sdrM* amplifications did not evolve significant resistance (**Fig. 6B**). While the *xerC*::Tn mutant populations in the initial tube evolutions evolved DLX resistance via mutations in the canonical targets and *sdrM*, the intergenic mutation between *B7H15_RS12090* and *B7H15_RS12095*, and less-prevalent *sdrM* amplifications (**Supp. Fig. 11**), none of the populations evolved significant DLX resistance in the secondary screen (**Fig. 6B**).

**Figure 6.**
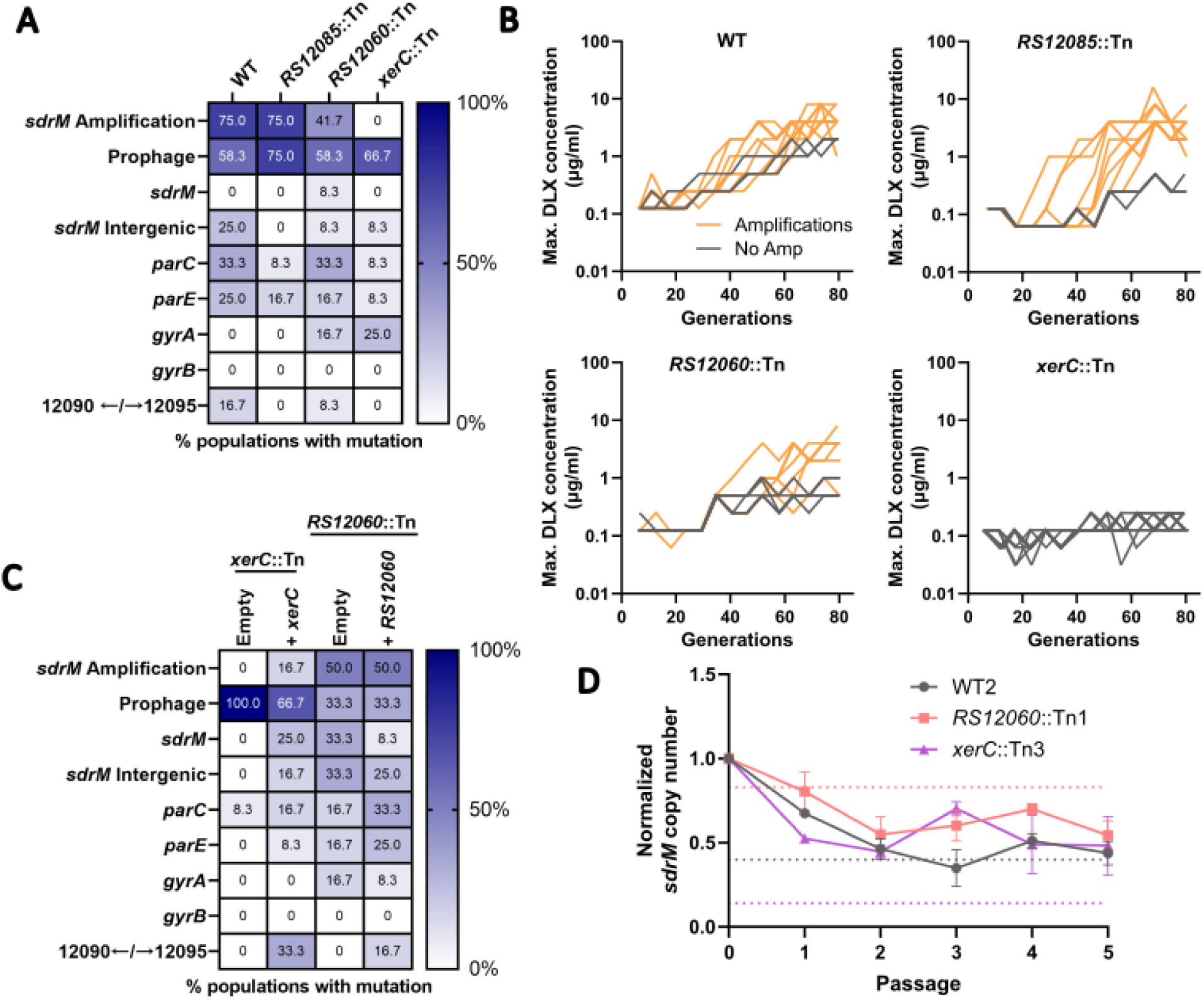
XerC is a novel determinant of amplification formation. **(A)** Percentage of 12 independently evolved populations of the WT, *RS12085*::Tn, *RS12060*::Tn, and *xerC*::Tn that showed an *sdrM* amplification, putative prophage induction, or mutations in the indicated genes. **(B)** Maximum DLX concentration that showed growth at each passage (shown as the respective number of generations) for the 12 independently evolved populations of WT, *RS12085*::Tn, *RS12060*::Tn, and *xerC*::Tn. Orange lines represent populations that evolved *sdrM* amplifications, while grey lines represent populations without *sdrM* amplifications. **(C)** Percentage of 12 independently evolved populations of the *xerC*::Tn and *RS12060*::Tn strains complemented with either an empty vector or one containing the respective gene, that showed an *sdrM* amplification, putative prophage induction, or mutations in the indicated genes. **(D)** Normalized *sdrM* copy number (normalized to the copy number in the initial culture) upon extended passaging without DLX for an evolved population each of WT, *RS12060*::Tn, and *xerC*::Tn containing an *sdrM* amplification. Dotted lines denote the absolute copy number for the respective samples. Data shown are the mean ± standard deviation of two independent passaging experiments.

The *B7H15_RS12060*::Tn and *xerC*::Tn mutants have transposon insertions in the respective genes, but the insertions could possibly cause polar effects, especially for *xerC*, which is the first gene in an operon that contains *hslUV*, encoding a heat-shock responsive quality control protease^43^, and *codY*, encoding a master regulator of metabolism and virulence^44^. To test whether *xerC* and *B7H15_RS12060* were involved in amplification formation, we complemented the corresponding genes back on a plasmid and repeated the deep well evolution (**Supp. Table 7**). We observed no change in the frequency of amplification formation upon complementation of *B7H15_RS12060*, but an increased frequency upon *xerC* complementation, demonstrating that *xerC* is a determinant of amplification formation (**Fig. 6C**). Finally, we passaged an evolved population with amplifications from each mutant background (from the tube evolutions) in DLX-free media, and observed no significant difference in the rate of loss of amplifications between the three strains (**Fig. 6D**). However, while the absolute copy number in the WT and *B7H15_RS12060*::Tn backgrounds reached a value of 1 or less, indicating a complete loss of amplifications, the absolute copy number remained high in the *xerC*::Tn background, indicating a possible role of XerC in the maintenance of *sdrM* amplifications.

## DISCUSSION

Gene amplifications can lead to resistance against several distinct antibiotics in *S. aureus*^8^, but modulators of such amplifications in *S. aureus* have not been described. In this study we used adaptive evolution to investigate the effectors involved in the formation and maintenance of *sdrM* gene amplifications that lead to DLX resistance in MRSA. Similar to what has been shown in other bacteria, we determined that functional RecA was required for *sdrM* gene amplifications, and was also involved in their expansion and contraction (**Fig. 3B and 3C**). Stable amplifications in the *recA*::Tn background led to a growth defect (**Fig. 3D**), indicating that in the absence of the selective pressure of DLX, maintenance of *sdrM* amplifications has a fitness cost, at least without functional RecA. Further, we identified that XerC is a novel effector of gene amplifications in MRSA, as mutants lacking functional XerC showed reduced rates of amplification formation and evolution of DLX resistance. Finally, mutants in other genes and pathways required for amplifications in *E. coli* still showed *sdrM* amplifications indicating that *S. aureus* may have alternate or redundant mechanisms for the generation of amplifications.

In the *recA*::Tn mutant, the evolution of DLX resistance was achieved in two steps, likely due to the hypersensitivity of this mutant to DNA damage caused by DLX. The first step involved acquiring mutations in DNA repair to reach the intrinsic WT DLX resistance levels, while the second step comprised of mutations in the canonical targets DNA gyrase and topoisomerase IV, as well as in *sdrM* (**Fig. 1A**). A similar trend was seen in the *rexA*::Tn mutant (**Supp. Fig. 4**). This suggests that in a DNA damage hypersensitive strain, repairing DNA damage, including that induced by DLX, provides a greater fitness benefit compared to increased DLX efflux, and is thus the first adaptation that is selected for.

A common mutation seen in the early steps of the *recA*::Tn evolution, albeit after mutations in *rexB* and *recD2* (**Fig. 1A**), as well as in the *rexA*::Tn evolution, was the intergenic mutation that lay within the predicted LexA binding site between the *B7H15_RS12090* and *B7H15_RS12095* genes^25^. This mutation conferred increased DNA damage resistance in the *recA*::Tn background (**Fig. 2E**), indicating enhanced DNA repair in the absence of RecA. Three of the 16 previously reported SOS regulon genes in *S. aureus* are thought to be controlled by that LexA binding site^25^, and we see increased expression of those genes in the mutant (**Fig. 5E**). While the function of these genes in DNA repair has not been investigated, and bioinformatic annotations do not reveal an obvious connection to DNA repair, their regulation by the SOS response, and increased DNA damage resistance in a mutant that overexpresses these genes points to a role in DNA repair. This mutant also showed elevated expression of *sdrM*, which was immediately downstream of the SOS operon. Further work is necessary to determine whether *sdrM* is in the SOS regulon, and whether the transcriptional overexpression seen in this mutant is only due to disruption of LexA binding. In a previous study, a mutation in the same LexA-binding site was seen in isolates of the *S. aureus* COL strain that were selected for resistance against oxadiazoles, a new class of β-lactams^46^. SOS induction has been reported in *S. aureus*^47^ and other species^48^ upon exposure to β-lactams, indicating a role of this mutation and the mis-regulated genes in adaptation against diverse stresses.

In our evolved populations, we observed a negative association between the presence of *sdrM* amplifications, and the LexA-binding site mutation or mutations in the upstream intergenic region of *sdrM*, both of which may increase *sdrM* expression. Further, a mutant carrying the mutation in the LexA-binding site showed only a transient amplification in just one out of three populations upon selection for DLX resistance (**Fig. 5D**). Such transient amplifications were rare in other strains where most *sdrM* amplifications became fixed in the population once they arose (**Supp. Table 5**), indicating that *sdrM* amplifications may not have a significant selective advantage when *sdrM* is already overexpressed. Similarly, the *recA*::Tn intermediate isolate R1 also had this LexA binding-site mutation, and it is likely that RecA complementation did not lead to *sdrM* amplifications in two of the three evolved populations as the parental strain (R1) already expressed *sdrM* at a higher level, and *sdrM* amplifications did not provide a significant further fitness advantage. Thus, these *sdrM* overexpressing mutations represent an alternate route of DLX resistance to *sdrM* amplifications. Notably, both these routes of DLX resistance center around increased *sdrM* expression, underscoring the critical role of *sdrM* function in DLX resistance.

Of the 105 populations that were evolved for DLX resistance, 89 populations had amplifications, while only 13 populations had at least one of the *sdrM* upstream or LexA-binding site mutations (of which 3 also had an *sdrM* amplification). Thus, *sdrM* amplifications are selected for at a significantly higher frequency compared to the specific point mutations, consistent with what has been reported before for the relative frequencies of these two types of mutations^7^.

The varied amplified *sdrM* fragments in our evolved populations showed limited homology between the fragment ends. Similar lack of homology in amplified fragments has been reported in other species such as *Acinetobacter* sp., *E. coli*, and *Salmonella*, upon exposure to diverse selection pressures^18,19,49^. Almost all our amplifications had one fragment end nearly 4 kb downstream of *sdrM* in the tRNA-rRNA loci, while the other end was in a region starting immediately upstream of *sdrM*. We postulate there are two, non-exclusive underlying reasons for this observation. First, amplification ends in the tRNA-rRNA locus could have a higher selective advantage as amplified copies of this locus would lie upstream of *sdrM* and could lead to higher *sdrM* expression due to potential read-through transcription, similar to what has been seen in *Streptococcus pneumoniae*^50^. Second, the highly transcribed tRNA-rRNA clusters are thought to be hotspots for codirectional replication-transcription conflicts^51^, and any resulting DNA damage may lead to a higher prevalence of duplications ending in this cluster.

Almost all our evolved populations showed increased coverage of an intact prophage (**Fig. 5A, Supp. Fig. 8**), likely indicating prophage induction due to DNA damage^52^. This prophage encodes a homolog of RecT, that has been implicated in recombination without homology or between short homologous sequences^53,54^. However, we did not detect significant prophage induction in a *recT*::Tn mutant (**Fig. 5A**), but still see *sdrM* amplifications, indicating that while RecT likely plays a role in the induction of this prophage, *sdrM* amplifications can occur in the absence of RecT and prophage induction. Further, there were multiple populations, e.g. in the *recX*::Tn, *rarA*::Tn, and *ruvX*::Tn mutants (**Supp. Table 5**), where we saw prophage induction, but no *sdrM* amplifications, suggesting these two events are unlinked.

Out of the 34 transposon mutants we evolved, only *recA*::Tn and *xerC*::Tn were significantly impaired in the formation or maintenance of *sdrM* gene amplifications. DNA repair pathways in *S. aureus* have been only partly characterized, and the roles of several DNA repair proteins as well as redundancies present in various repair pathways have not been fully dissected^23,40^. The list of screened mutants included those with transposon insertions in genes involved in double-strand break repair, nucleotide excision repair, Holliday junction resolution, and cell division (**Supp. Table 3**)^23,40^. In *E. coli*, it has been shown that mutants in double-strand break repair (*recA*, *recB*, *ruvC*) are necessary for amplifications^19^, while in *Salmonella enterica* Typhimurium, individual mutations in *recB* and *recF* did not significantly alter the frequency of formation of duplications (the formation of amplifications was not tested), while a *recA* mutant and a *recB recF* double mutant showed significantly lower rates of duplications^18^. Our data show that unlike *E. coli*, a mutant in RexA, which is part of the RexAB helicase/nuclease that performs a similar function as RecBCD in double-strand break repair, still evolves amplifications. Given the requirement of RecA, this indicates that other helicase/nuclease combinations may function with RecA in homologous recombination during amplification formation. Further, while DNA polymerase I (PolA) is required for amplifications in *E. coli*^19,55^, the *S. aureus polA*::Tn mutant showed prevalent amplifications. Similar to *E. coli*^19^, mutations in DNA polymerases IV and V (DinB and UmuC) still allowed for amplifications, indicating that in *S. aureus* other polymerases may play a role in amplifications.

We found that the lack of a functional tyrosine recombinase XerC significantly reduced the frequency of *sdrM* amplification formation. XerC is a widely conserved protein involved in the segregation of daughter chromosomes during cell division, and functions with the related recombinase XerD, and the DNA translocase FtsK^42,56^. In *S. aureus*, the *xerC*::Tn mutant is deficient in the induction of the SOS response, and has increased sensitivity to DNA gyrase inhibiting and cell-wall synthesis targeting antibiotics^58^. However, any role of XerC in the formation or selection of gene amplifications remains unknown. In *S. aureus*, XerD is an essential protein, unlike in other species, suggesting that it may play additional functions^56^. In contrast to the *recA*::Tn mutant, in our tube evolutions, one out of the three *xerC*::Tn populations evolved *sdrM* amplifications albeit in the later passages (**Supp. Table 10**). Future work will focus on the mechanism by which XerC promotes amplification formation.

Gene amplifications can lead to rapid evolution of resistance against many classes of antibiotics, including ones that have multiple targets in the cell^8,16^. Pathways required for the formation of such amplifications can thus be putative targets in combination with antibiotics, to lower the frequency of resistance evolution. Our data suggest that apart from RecA and XerC, genes with redundant function may play a role in the formation of gene amplifications in *S. aureus*, and that such genes and pathways may differ between bacterial species. Combinatorial mutagenesis, and genome-wide screens may be essential to identify additional determinants of amplification formation.

## METHODS

### Strains and growth conditions

All strains and plasmids used in this study are listed in **Supplementary Table 8**.

For experiments in liquid media, bacteria were grown at 37°C, shaking at 300 rpm in modified M63 media (13.6 g/L KH2PO4, 2g/L (NH4)2SO4,0.4 μM ferric citrate, 1mM MgSO4; pH adjusted to 7.0 with KOH) supplemented with 0.3% glucose, 0.1 μg/ml biotin, 2 μg/ml nicotinic acid, 1× Supplement EZ (Teknova) and 1× ACGU solution (Teknova)^16^. For experiments with deep well plates, the plates were incubated in a Titramax 1000 (Heidolph) incubator at 37°C shaking at 900 rpm. To construct and mutate plasmids and strains, cells were grown in LB liquid medium (10 g/L bacto-tryptone, 5 g/L yeast extract, 10 g/L NaCl) at 25°C, 30°C or 37°C as indicated, shaking at 300 rpm, or on LB plates (15 g/L agar) supplemented with the appropriate antibiotics (10 μg/ml chloramphenicol, 50 μg/ml kanamycin, 10 μg/ml trimethoprim, 10 μg/ml erythromycin) or 0.4% para-chlorophenylalanine (PCPA).

### Minimum inhibitory concentration measurements

MICs were tested as described previously in M63 media^16^. Briefly, a 96-well flat clear bottom plate (Corning) was used to prepare two-fold serial dilutions with eight concentrations of DLX. Cells grown overnight in M63 were diluted 1:5000 in fresh M63 and added at a 1:1 ratio to the 96-well plate with the DLX dilutions such that the final dilution of the cells was 1:10000. Cells were grown for 24 hours in a Titramax 1000 (Heidolph) incubator at 37 °C with shaking at 900 rpm. The *recA*::Tn, 1.7a *recA*::Tn, *rexA*::Tn, and *xerC*::Tn strains were grown for an additional 24 hours to compensate for their slower growth. Following the growth, OD_600_ was measured using a microplate reader (Biotek Synergy H1) and the MIC values were determined by fitting the OD_600_ vs. DLX concentration to a modified Gompertz function^59^.

### Growth curves and analysis

Cells were grown overnight in tubes with 2 ml of M63, diluted 1:100 in 200μl M63 and added to a COSTAR clear bottom 96-well plate. OD_600_ was measured using a microplate reader (Biotek Synergy H1) every 30min for 24hrs, while shaking at 800 rpm at 37 °C. The growth curves were analyzed with the R package Growthcurver to determine growth rate, doubling time and the time to reach the maximum growth rate^60^.

### Experimental evolution of DLX resistance in tubes

Evolution of *S. aureus* JE2 cells was carried out as described previously^16^. Briefly, for most strains, three independent colonies were picked and cultured in 2ml of M63. After overnight growth, cells were transferred to new tubes containing 2 mL M63 with DLX at a concentration ∼0.5x the respective MIC. After 24h growth, 40 μl cells were transferred to tubes containing twice the concentration of DLX as the prior passage in 2 ml M63 and this was repeated up to a DLX concentration of 2 μg/ml. If sufficient growth was not apparent after 24 hours, cells were given an additional 24-48 hours to grow. If growth was still not seen, the evolution was reset to the previous passage and the DLX increment was adjusted depending on the growth. If no gene amplifications were detected in the final passage (2 μg/ml), evolutions were continued similarly and carried out until a terminal concentration of 8 or 16 μg/ml.

Given that the *recA*::Tn and *rexA*::Tn mutants did not evolve canonical resistance mutations or *sdrM* amplifications until the evolving populations reached an MIC near the WT MIC, we decided on a slightly modified strategy to evolve other mutants that had similarly low DLX MICs. For these mutants (*xerC*::Tn and *recG*::Tn) which had very low MICs (<10-fold the WT MIC), a single population was evolved until the DLX concentration in the evolution approached the JE2 WT MIC (∼0.22 μg/ml). For the next passage, cells were cultured with a dilution of 1:50 into three independent tubes containing twice the previous DLX concentration, and subsequently these three populations were independently evolved as mentioned above. For the *recA*::Tn and *rexA*::Tn mutants, we sequenced intermittent passages beginning from those that grew in a DLX concentration one-tenth of the WT MIC, i.e. 0.025 µg/mL DLX. For the other tube evolutions, we sequenced the passage that grew at 2 µg/mL, and if subsequent evolutions were performed, we sequenced intermittent passages, as well as the terminal one.

### Laboratory deep well evolution of DLX resistance

Mutants that needed to be tested further for amplification frequency were streaked out on selective plates, and 12 independent colonies per strain were cultured in 400 μl of M63 in deep well plates (Nunc™ 96 DeepWell™ Polystyrene Plates) for 24hrs with shaking at 37°C. For the *xerC*::Tn mutant, the original evolved population (prior to splitting the evolution into three independent populations as described above), that had an MIC similar to the WT, was streaked out on a plate. A single colony was re-streaked out, and 12 colonies from this plate were cultured to start the 12 independent populations. 4 μl cells were then transferred to a fresh deep well plate containing 400 μl of M63 with 8 serially diluted DLX concentrations with the maximum DLX concentration set at 1-2μg/ml. This meant that each independent strain was cultured in all the 8 DLX concentrations. For the following passages, 8μl of cells in the well at the highest DLX concentration that showed growth (defined by OD_600_ > 0.400 as measured in a Biotek Synergy H1 microplate reader) was transferred into a new deep well plate containing the same DLX concentration profile and grown for 24hrs. The maximum DLX concentration for the subsequent passages was increased if there was growth in the 2^nd^ well of the previous passage. The evolution was continued for 14 passages in total and the genomic DNA of the terminal passage was sent out for whole genome sequencing.

### Whole genome sequencing

Genomic DNA was extracted from cultures using the Qiagen DNeasy Blood and Tissue kit. Sequencing libraries were prepared using the Illumina Nextera XT DNA Library Preparation kit and sequenced (75 cycle or 100 cycle) on the Illumina NextSeq 550 (75 cycle) or the NextSeq 2000 (100 cycle) instruments to obtain single-end reads. The FASTQ files were processed by removing adapters and trimming using fastp v0.23.2. Sequences were aligned, variants called, and novel junctions identified using breseq v0.37.17. The mutations were marked as present in the respective figures (**Figs. 1A, 1D, 5A, 5B, 5D**, and **Supp. Figs. 5 and 11**) only if at least 30% of the population contained the particular mutation. Copy number variation in the efflux pump *sdrM* was identified as described previously^16^. Briefly, in breseq we used the BAM2COV command to determine the read coverage depth across the whole genome. The *sdrM* copy number was determined by dividing the average coverage depth of *sdrM* with the average coverage depth of the whole genome (both normalized to length). The threshold used to define an *sdrM* amplification was the presence of a novel junction around *sdrM* and an *sdrM* copy number greater than 1.3x the WT *sdrM* copy number.

### Construction of allele replacement and overexpression strains in *S. aureus* JE2

Construction of the allele replacement mutants and overexpression strains in JE2 was done as described previously^16^. Briefly, for the allele replacement, the new allele with flanking homology was cloned into the pIMAY* plasmid^61^, both of which had been PCR amplified. The plasmid was then electroporated into *S. aureus* JE2. Strains carrying the integrated plasmid were selected for at 37°C (the selective temperature), and the plasmids were subsequently excised via PCPA counter selection, and plasmid loss was confirmed via testing for chloramphenicol sensitivity. The final mutant was confirmed by flanking PCR and Sanger sequencing.

To construct the overexpression strains, the appropriate genes with their native promoters were cloned into the pKK30 vector and electroporated into *S. aureus* JE2. Primers used for constructing all mutants and overexpression strains are listed in **Supp. Table 9**.

### Mutagenesis of *lexA* gene

The Agilent QuikChange Lightning Site-Directed Mutagenesis Kit was used as per the manufacturer’s instructions to generate the *lexA* mutant alleles. Mutagenic primers were created utilizing the Agilent web-based QuikChange Primer Design tool and are listed in **Supp. Table 9**.

### Phage transduction of *recA::*Tn into 1.7a

The donor *recA*::Tn strain was grown overnight in LB, diluted 100-fold in fresh LB the following day, and grown at 30°C for 1.5-2 hrs. Then, 1 ml of 10 mg/ml CaCl_2_ and 10 μl of 10^10^ bacteriophage 85 (phi85) suspension was added to this culture which was incubated at room temperature for 30 min without shaking and then rotated slowly (80RPM) at 37°C for 5–6 hrs. The culture was then kept static at 4°C overnight. The next day, the culture was filtered using a 0.45 μm filter and the phage-containing filtrate was stored at 4°C.

To transduce the *recA*::Tn mutation to the appropriate JE2 background, we grew 2 ml of the recipient cells overnight in LB, and collected the cell pellets the following day by centrifugation. Cell pellets were resuspended in 300 μl of LB broth with 5 mM CaCl_2_ and 700 μl of the *recA*::Tn phi85 phages were added. This mixture was incubated at 37°C without shaking for 20 min. Cells were then washed multiple times with 1ml of 40mM sodium citrate and finally plated on LB + erythromycin plates containing 200 μM sodium citrate. The *recA*::Tn transposon insertion as confirmed via PCR.

### Quantitative PCR (qPCR)

For quantifying *sdrM* copy number, qPCR was done as described previously^16^. Briefly, gDNA samples diluted to 10 ng/µl were mixed with appropriate qPCR primers and Applied Biosystems Power SYBR Green PCR Master Mix (Thermo Scientific) in a Microamp EnduraPlate Optical 384 Well Clear Reaction Plate (Thermo Fisher Scientific) and the qPCR was carried out in a QuantStudio 5 real-time PCR machine (Thermo Fisher Scientific). We used *rpoC* as the control housekeeping gene. To determine the limit of detection for the junctions, gDNA of the evolved population at 10 ng/µl was serially diluted 10-fold in 9μl of genomic DNA from the parental WT strain (10 ng/µl) seven consecutive times. Then the qPCR was carried out as described above.

### Reverse transcription quantitative PCR (RT-qPCR)

To determine the expression level of the genes neighboring the *lexA* binding site including *sdrM*, cells grown overnight in tubes with 2 ml M63 were diluted 1:100 in 10 ml of M63 in a flask and grown for an additional 3-4 hours to an OD_600_ of 0.3-0.4. For the *sdrM* allele-replacement strains, we similarly grew the cells to an OD_600_ of ∼0.5 and also took a sample of overnight grown cells. 2x volume of RNAprotect Bacteria Reagent (QIAGEN) was added to the culture and after 10 min incubation at room temperature, cells were centrifuged at 4000 rpm, the supernatant removed, and the pellets stored at-80 °C. RNA was extracted using the NORGEN BIOTEK Total RNA purification kit and genomic DNA was removed with the TURBO DNA-free™ Kit (Invitrogen). Absence of genomic DNA was confirmed via PCR. cDNA was synthesized utilizing the Superscript III Reverse Transcriptase (Thermo Fisher Scientific) with random primers. The subsequent qPCR was carried out as described in the previous section.

### TOPO TA cloning

Genomic DNA used as a template for the PCR reactions was concentrated to ∼200 ng/µl and 2 µl of it was added to each 23 µl PCR reaction. PCR reactions were performed using the OneTaq or LongAmp polymerases, purified with the Zymo Clean and Concentrator-5 kit, and cloned into the pCR™4-TOPO TA plasmid using the TOPO TA cloning kit for Sequencing, with One Shot™ TOP10 Chemically Competent *E. coli* (Invitrogen). Blue-white selection was used to identify colonies carrying plasmids with inserts by supplementing the LB + Kanamycin plates with 40 μg/ml of X-gal (Invitrogen). Plasmids were extracted using the Qiaprep Spin Miniprep kit (Qiagen), and sent for whole plasmid sequencing to identify the inserted fragments.

### Determination of amplification stability

For the amplification stability experiment with 1.7a (*recA*::Tn), cells were inoculated into M63 from the corresponding frozen stock and grown overnight. For the amplification loss experiment, which was done for five passages, cells were grown from frozen stocks into M63 containing DLX at ∼0.25x the corresponding MICs, in order to stabilize the amplifications prior to the treatment free passaging. The cultures were then diluted 1:1000 in 2 ml of M63 (with or without DLX) and grown for 24 hrs. Genomic DNA was extracted from the remaining culture. This procedure was repeated for the appropriate numbers of passages, and WGS was performed on the genomic DNA from each passage. Copy number for *sdrM* was determined as described above.

### TUNEL assay

We followed the terminal deoxynucleotide transferase dUTP nick end labeling (TUNEL) assay as described previously for bacteria^62^. Briefly, cells were grown overnight in 2 ml of M63, diluted 1:50 in 15 ml of M63, and grown to an OD_600_ of 0.3-0.4. Next, 2 ml of cells were transferred to three falcon tubes, and two of the tubes were treated with the appropriate concentration of DLX or MMC respectively. The remaining tube was kept as a treatment free control, and all three tubes were grown for an additional 3 hrs. From these cultures, 1 ml was transferred to microcentrifuge tubes, washed once with ice cold 1x Phosphate buffered saline (PBS,1X, Thermo Scientific), and fixed by treatment with 1ml of 4% Paraformaldehyde (PFA) on ice for 30 min. Cells were then washed with ice cold PBS once, and resuspended in 250 μl of PBS and 1ml of ice cold 70% ethanol, and incubated 12-16 hours at-20°C.

After incubation, the tubes were centrifuged, the supernatants removed, and the cells labeled with the Apo-Direct Kit (BD Bioscience). Fluorescence output was analyzed using a Apogee MicroPLUS flow cytometer (ApogeeFlow Systems Inc), using excitation and emission wavelengths of 488 nm and 515 nm respectively. A region of interest (ROI) was drawn around the WT cell events for medium vs large light scatter angle, and this ROI was used as a gate for subsequent analysis of other strains. The proportion of the population with DNA damage was quantified as the Overton positive percentage using FlowJo (v10.1)^63^.

### Statistics

Statistical comparison of datasets was performed using GraphPad Prism 9. Tests used and significance thresholds are mentioned in the respective figure legends.

### Data Availability

The whole genome sequencing (WGS) data have been deposited at NCBI Short Read Archive (SRA) under the bioproject PRJNA1242835.

## ACKNOWLEDGEMENTS

We would like to acknowledge the Center for Cancer Research (CCR) Genomics Core for whole-genome sequencing. This work used the computational resources of the NIH High Performance Computing Biowulf Cluster (http://hpc.nih.gov). We thank Susan Gottesman, Gisela Storz, and Anthony Martini for comments on the manuscript, and members of the Khare, Gottesman, and Ramamurthi labs for helpful discussion and feedback. This work was supported by the Intramural Research Program of the NIH, National Cancer Institute, Center for Cancer Research.

