## Supplemental Figures and Supplemental Tables 8 and 9 for "Genetic determinants of gene amplifications alter frequency and evolutionary trajectory of antibiotic resistance in *Staphylococcus aureus*"

### SUPPLEMENTARY FIGURES

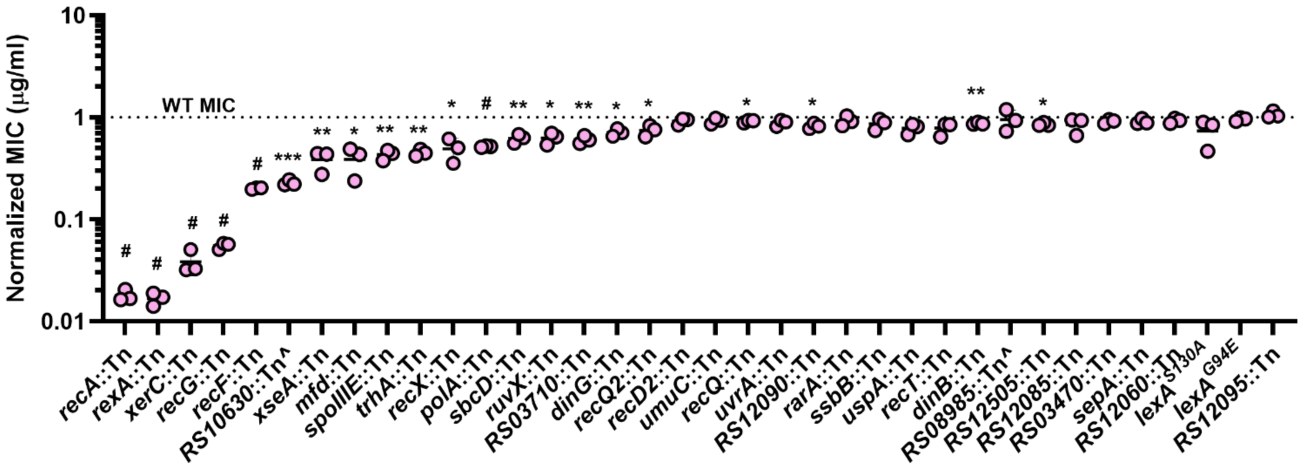

**Supp. Fig. 1. Selected mutants show a wide range of DLX sensitivities.** DLX MICs of WT and the indicated mutants were measured, and the mutant MICs were normalized to the WT MIC. Data shown are the mean ± standard deviation for three biological replicates. Significance is shown for comparison to a value of 1, as tested by one-sample *t*-tests (* $p < 0.05$, ** $p < 0.01$, *** $p < 0.001$, # $p < 0.0001$).

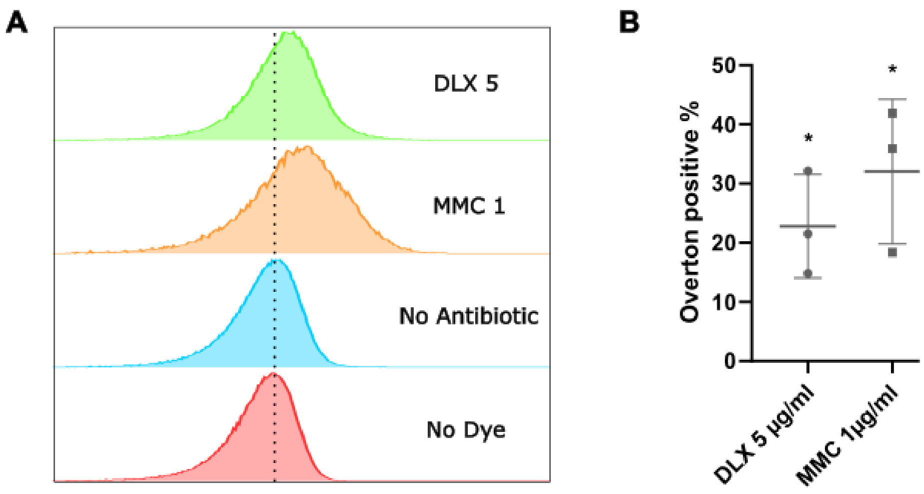

**Supp. Fig. 2. DLX causes DNA damage. (A)** Representative fluorescence from TUNEL staining of WT cells treated with no antibiotic, 1 µg/mL MMC, or 5 µg/mL DLX, or a no dye control. **(B)** The Overton positive percentage (representing the percentage of the population that has increased fluorescence compared to the control no antibiotic population) for the DLX and MMC treated samples. Significance is shown for comparison to a value of 0, as tested by one-sample *t*-tests (* $p < 0.05$).

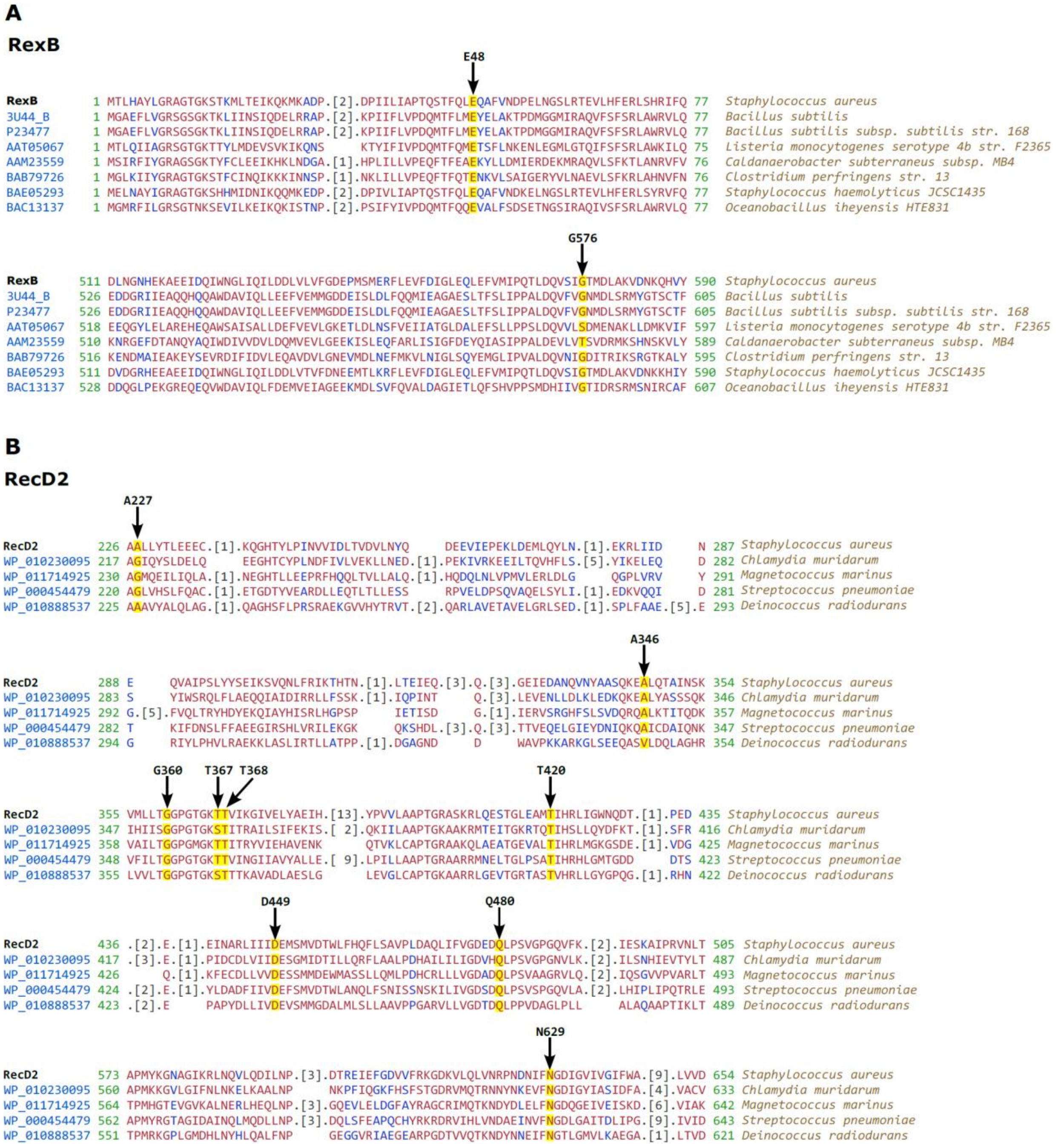

**Supp. Fig. 3. Mutations seen in RexB and RecD2 are in relatively conserved residues. (A) and (B)** Multiple sequence alignments of **(A)** RexB and **(B)** RecD2 from the Conserved Domain Database[1,2], where residues in red are highly conserved, and those in blue are not. Arrows denote the position of mutations, which are also highlighted, seen in the *recA*::Tn evolved populations.

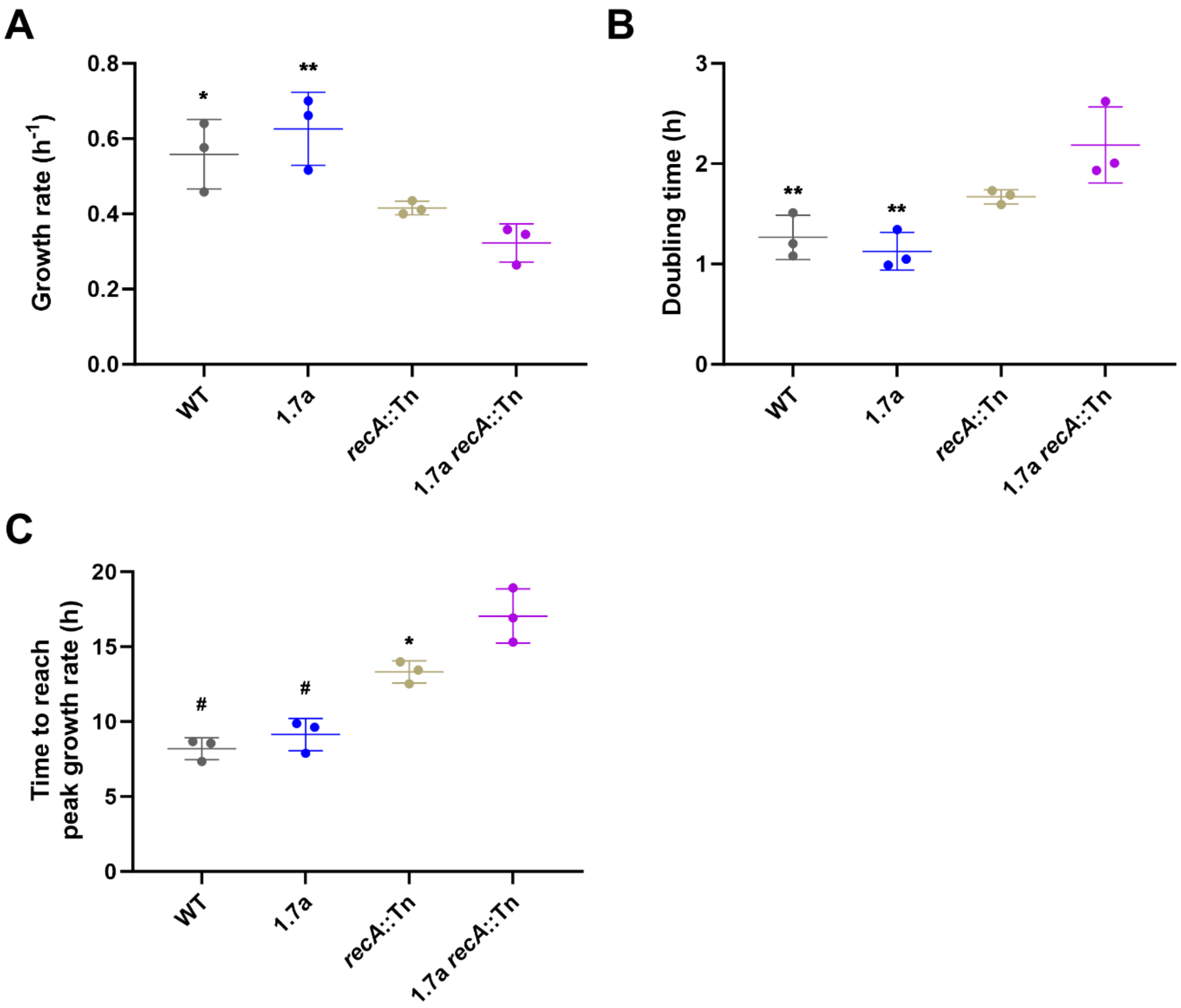

**Supp. Fig. 4. Amplifications of *sdrM* have a fitness cost. (A)** Growth rates, **(B)** doubling times, and **(C)** time to reach the peak growth rate are shown for the WT, and 1.7a, and *recA*::Tn mutants in both backgrounds. These parameters were obtained from growth curves shown in **Figure 3D** using Growthcurver[3] for fitting and analysis. Data shown are the means ± standard deviation for three biological replicates. Significance is shown for comparison to the 1.7a *recA*::Tn strain, as tested by a one-way ANOVA with Dunnett's test for multiple comparisons (* $p < 0.05$, ** $p < 0.01$, # $p < 0.0001$).

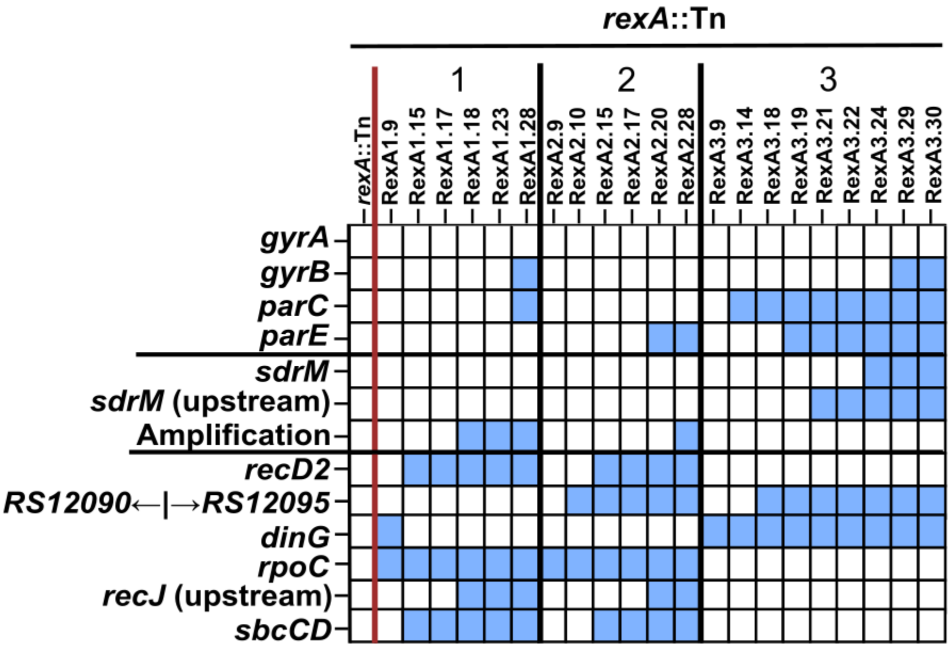

**Supp. Fig. 5. The *rexA*::Tn mutant evolves DLX resistance initially via DNA repair-associated mutations, and later via mutations in the canonical targets, *sdrM*, mutations increasing *sdrM* expression, and *sdrM* amplifications.** Common mutations seen in three independently evolved populations of the *rexA*::Tn mutant. Presence of mutations in the corresponding genes is indicated by a blue square. For each population, passages are shown from left to right in chronological order.

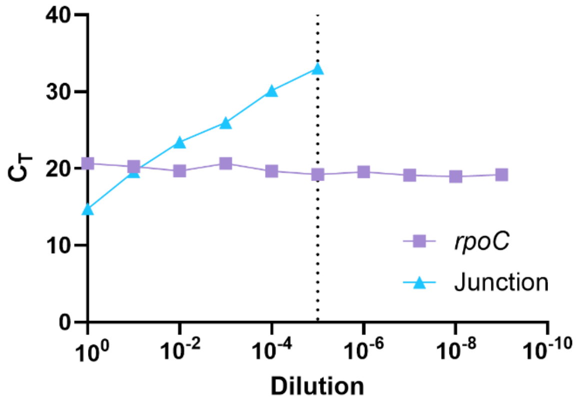

**Supp. Fig. 6. Novel junctions corresponding to the *sdrM* amplifications can be detected at a $10^{-4}$ – $10^{-5}$ dilution.** qPCR $C_T$ values for *rpoC* and the respective novel junction present in two different WT evolved DLX resistant populations, measured in a 10-fold dilution series where genomic DNA from the evolved population was diluted in genomic DNA from the WT strain.

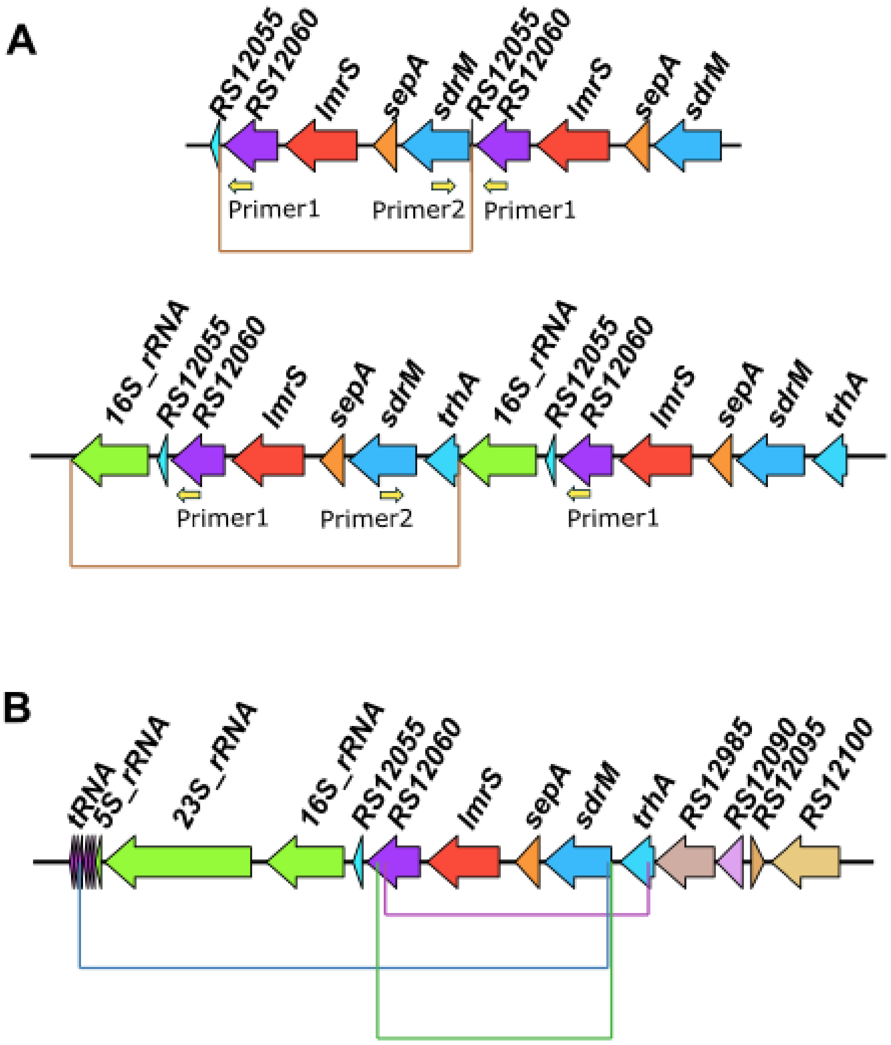

**Supp. Fig. 7. Rare novel junctions around *sdrM* may be present in the unselected WT strain. (A)** Inverse-PCR like strategy to detect novel junctions present around *sdrM*. Shown are two hypothetical amplifications (outlined by the brown lines). We designed outward facing primers (Primer 1 and Primer 2) binding to *sdrM* and *B7H15_RS12060*. If a duplication or amplification is present in that region, the novel junction should get amplified by the PCR primers. **(B)** Three putative duplications or amplifications identified in the WT strain using the inverse PCR strategy.

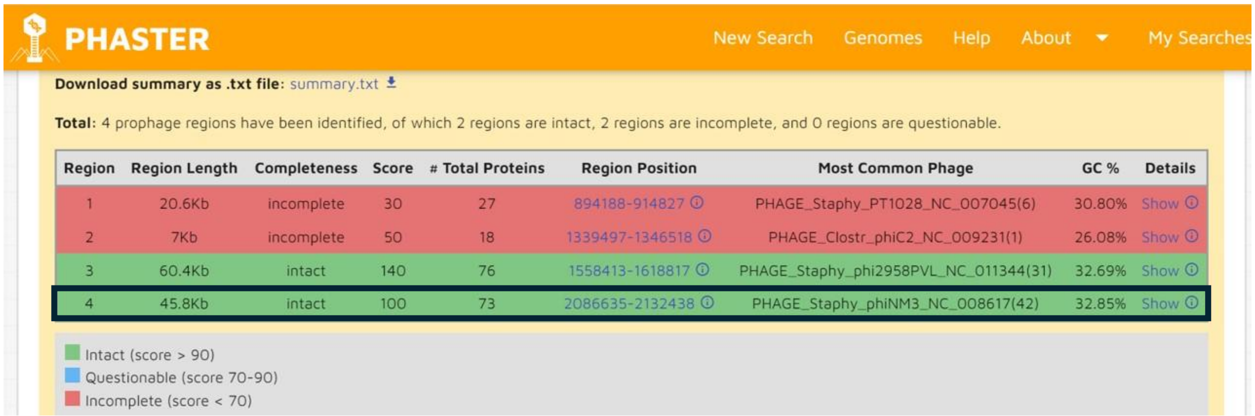

PHASTER New Search Genomes Help About My Searches

**Download summary as .txt file:** summary.txt

**Total:** 4 prophage regions have been identified, of which 2 regions are intact, 2 regions are incomplete, and 0 regions are questionable.

| Region | Region Length | Completeness | Score | # Total Proteins | Region Position | Most Common Phage | GC % | Details |
|---|---|---|---|---|---|---|---|---|
| 1 | 20.6Kb | incomplete | 30 | 27 | 894188-914827 | PHAGE_Staphy_PT1028_NC_007045(6) | 30.80% | Show |
| 2 | 7Kb | incomplete | 50 | 18 | 1339497-1346518 | PHAGE_Clostr_phiC2_NC_009231(1) | 26.08% | Show |
| 3 | 60.4Kb | intact | 140 | 76 | 1558413-1618817 | PHAGE_Staphy_phi2958PVL_NC_011344(31) | 32.69% | Show |
| 4 | 45.8Kb | intact | 100 | 73 | 2086635-2132438 | PHAGE_Staphy_phiNM3_NC_008617(42) | 32.85% | Show |

Intact (score > 90)
Questionable (score 70-90)
Incomplete (score < 70)

**Supp. Fig. 8. The second region with increased coverage in several DLX resistant evolved populations encodes for a putative intact prophage.** The JE2 genome was analyzed using PHASTER[4] to identify prophage regions. Region 4 (outlined in black) encodes an intact prophage whose coordinates match the region that shows increased sequencing coverage depth in the evolved DLX resistant populations.

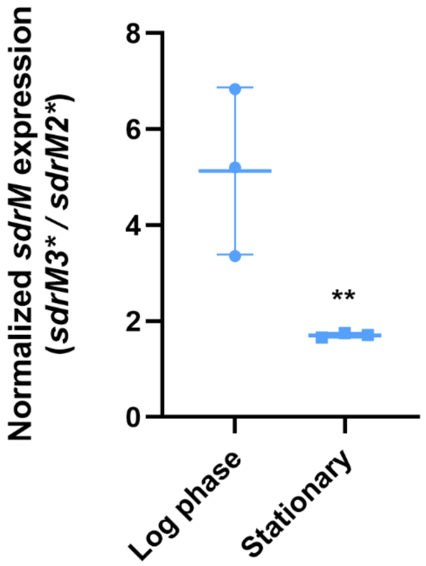

**Supp. Fig. 9. An intergenic mutation upstream of *sdrM* leads to increased *sdrM* expression.** Expression of *sdrM* was measured in log phase ($OD_{600}$ = 0.5) and stationary phase (overnight cultures) of the previously reported *sdrM2** and *sdrM3** allele-replacement mutants[5]. The *sdrM2** mutant has the A268S coding sequence mutation, while the *sdrM3** mutant has the A268S mutation as well as a C to G change at position -164 (upstream of *sdrM*). Data shown are the means ± standard deviation for three biological replicates. Significance is shown for comparison to 1, as tested by a one-sample *t*-test (** $p < 0.01$).

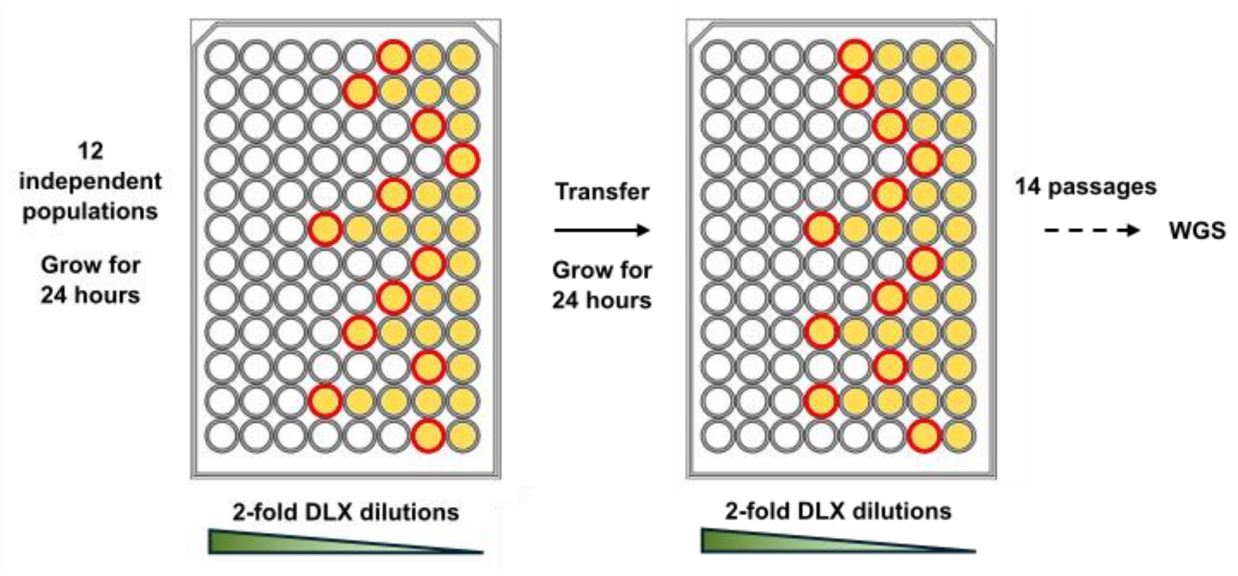

**Supp. Fig. 10. Scheme for deep well evolutions.** 12 independent populations of the strain to be evolved were inoculated in a 2-fold dilution series of DLX and grown for 24 hours. Populations from the well with the highest DLX concentration that showed growth were transferred to the next passage. Wells showing growth are denoted in yellow, and of those, the ones with the highest DLX concentration are marked with the red circles. Populations from these red-circled wells were propagated to the next passage. After 14 passages, the terminal passages from the wells with the highest DLX concentration that showed growth were sent for WGS.

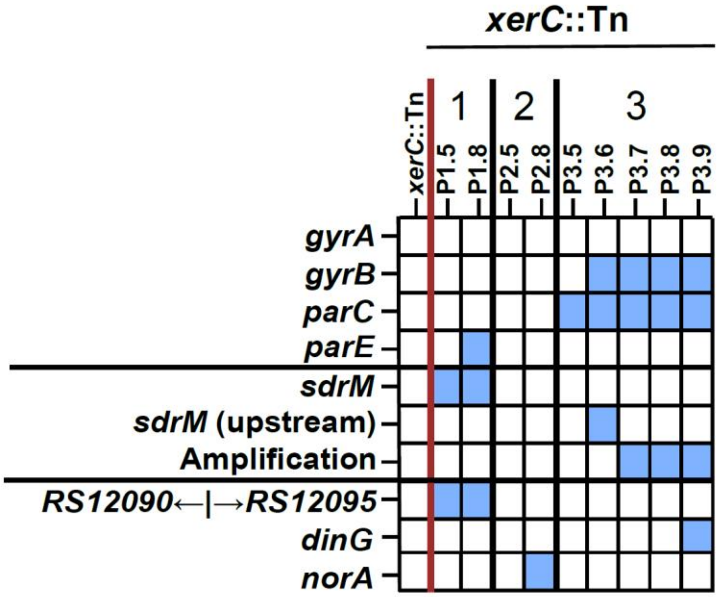

**Supp. Fig. 11. The *xerC*::Tn mutant evolves DLX resistance via mutations in the canonical targets, *sdrM*, mutations increasing *sdrM* expression, and less prevalent *sdrM* amplifications.** Common mutations seen in three independently evolved populations of the *xerC*::Tn mutant. Presence of mutations in the corresponding genes is indicated by a blue square. For each population, passages are shown from left to right in chronological order.

### SUPPLEMENTARY TABLES

**Supp. Table 1. Mutations seen in the *recA*::Tn evolved populations.**

**Supp. Table 2. Mutations seen in the evolved populations of R1 complemented with either an empty pKK30 plasmid or one carrying *recA*.**

**Supp. Table 3. List of strains evolved for DLX resistance.**

**Supp. Table 4. The distinct amplification fragments observed in the evolved populations from the tube evolutions of transposon and allele-replacement mutants.**

**Supp. Table 5. Mutations seen in the evolved populations from the tube evolutions of transposon and allele-replacement mutants.**

**Supp. Table 6. Mutations seen in the evolved populations of the allele-replacement strain with an intergenic mutation between *B7H15_RS12090* and *B7H15_RS12095*.**

**Supp. Table 7. Mutations seen in the evolved populations from the deep-well evolutions of transposon and complemented mutants.**

**Supplementary Table 8. Strains and plasmids used in this study.**

| Name | Description | Source |
|---|---|---|
| **Strains** | | |
| ***S. aureus*** | | |
| JE2 | *Staphylococcus aureus* subsp. *aureus* USA300_FPR3757 (CA-MRSA)-JE2 | |
| RN4220 | Restriction modification deficient *S. aureus*; used to shuttle pKK30 derivatives into JE2 | |
| SB514 | 1.7a (evolved DLX resistant mutant strain) | [5] |
| SB616 | JE2 *lexA*$^{S130A}$ | This study |
| SB617 | JE2 *lexA*$^{G94E}$ | This study |
| NE805 | JE2 *recA*::Tn | [6] |
| NE331 | JE2 *rexA*::Tn | [6] |
| NE1427 | JE2 *recD2*::Tn | [6] |
| NE959 | JE2 *sbcD*::Tn | [6] |
| NE324 | JE2 *recX*::Tn | [6] |
| NE1673 | JE2 *B7H15_RS12095*::Tn | [6] |
| NE883 | JE2 *xerC*::Tn | [6] |
| NE445 | JE2 *umuC*::Tn | [6] |
| NE1344 | JE2 *recG*::Tn | [6] |
| NE346 | JE2 *dinG*::Tn | [6] |
| NE972 | JE2 *recQ*::Tn | [6] |
| NE1146 | JE2 *B7H15_RS12090*::Tn | [6] |
| NE145 | JE2 *uvrA*::Tn | [6] |
| NE1794 | JE2 *ruvX*::Tn | [6] |
| NE555 | JE2 *recF*::Tn | [6] |
| NE711 | JE2 *B7H15_RS09290*::Tn | [6] |
| NE341 | JE2 *rarA*::Tn | [6] |
| NE458 | JE2 *xseA*::Tn | [6] |
| NE1679 | JE2 *ssbB*::Tn | [6] |
| NE1830 | JE2 *recT*::Tn | [6] |
| NE1528 | JE2 *recQ2*::Tn | [6] |
| NE1866 | JE2 *dinB*::Tn | [6] |
| NE188 | JE2 *mfd*::Tn | [6] |
| NE97 | JE2 *B7H15_RS12085*::Tn | [6] |
| NE1760 | JE2 *B7H15_RS08985*::Tn | [6] |
| NE993 | JE2 *trhA*::Tn | [6] |
| NE1689 | JE2 *B7H15_RS10630*::Tn | [6] |
| NE1616 | JE2 *sepA*::Tn | [6] |
| NE100 | JE2 *B7H15_RS03470*::Tn | [6] |
| NE348 | JE2 *B7H15_RS06660*::Tn | [6] |
| NE1529 | JE2 *B7H15_RS12060*::Tn | [6] |
| NE1915 | JE2 *B7H15_RS03710*::Tn | [6] |
| NE22 | JE2 *polA*::Tn | [6] |
| NE152 | JE2 *B7H15_RS12505*::Tn | [6] |
| SB618 | Evolved *recA*::Tn isolate RecA1.P14a (R1) | This study |

| | | |
|---|---|---|
| SB619 | Evolved *recA*::Tn isolate RecA2.P13a (R2) | This study |
| SB620 | Evolved *recA*::Tn isolate RecA3.P14a (R3) | This study |
| SB622 | 1.7a *recA*::Tn | This study |
| SB482 | JE2 *recD2*$^{A227E}$ | This study |
| SB483 | JE2 *rexB*$^{G576D}$ | This study |
| SB484 | JE2 *B7H15_RS12090* ← / → *B7H15_RS12095*, at position 2,306,688, G→A | This study |
| SB486 | *recA*::Tn *recD2*$^{A227E}$ | This study |
| SB487 | *recA*::Tn *rexB*$^{G576D}$ | This study |
| SB489 | *recA*::Tn B7H15_12090 ← / → B7H15_12095, at position 2,306,688, G→A | This study |
| SB621 | Evolved DLX-resistant *xerC*::Tn mutant XC5_7 | This study |
| ***E. coli*** | | |
| DH5α-λpir | DH5α lysogenized with λpir; host for pKK30 | [7] |
| IM08B | For plasmid transfer into *S. aureus* | [8] |
| **Plasmids** | | |
| pIMAY* | *S. aureus* allelic exchange plasmid; Cm$^R$ | [9] |
| pKK30 | Expression vector for *S. aureus*; Tmp$^R$ | [10] |
| pSB627 | pIMAY*-*recD2*$^{A227E}$ | This study |
| pSB626 | pIMAY*-*rexB*$^{G576D}$ | This study |
| pSB628 | pIMAY*- *RS12090* ←/→ *RS12095* | This study |
| pSB623 | pIMAY*-*lexA* | This study |
| pSB624 | pIMAY*-*lexA*$^{S130A}$ | This study |
| pSB625 | pIMAY*-*lexA*$^{G94E}$ | This study |
| pSB469 | pKK30-*recA* | This study |
| pSB598 | pKK30-*xerC* | This study |
| pSB600 | pKK30- *RS12060* | This study |

Cm$^R$: chloramphenicol resistant; Tmp$^R$: trimethoprim resistant

**Supplementary Table 9. Primers used in this study.**

| Primer Description | Primer Sequence* |
|---|---|
| Amplify *rexB* for pIMAY* F | TATCGATAAGCTTGATATCGAATTTAAACATAATTTCAATCCTGAAAATAC |
| Amplify *rexB* for pIMAY* R | TGGAGCTCCACCGCGGTGGCCTCAACTAGTACAGCTGTTTTAC |
| Check pIMAY*-*rexB*$^{G576D}$ integration F | ATACAACATCGTTTGTCGGTTT |
| Check pIMAY*-*rexB*$^{G576D}$ integration R | TTGCGCGTCAGTCCAAAT |
| Sanger sequencing for *rexB*$^{G576D}$ | GGCGTAAAGCGCATAAACTG |
| Amplify *recD2* for pIMAY* F | TATCGATAAGCTTGATATCGAAAAATTGATGACACAGATAGTC |
| Amplify *recD2* for pIMAY* R | TGGAGCTCCACCGCGGTGGCAACTCAACTTAAACATGTATTAAC |
| Check pIMAY*-*recD2*$^{A227E}$ integration F | CGGTACCACCCTAGTTATAAATGC |
| Check pIMAY*- *recD2*$^{A227E}$ integration R | ATGCAAGATACAATTTGGGCTTAG |
| Sanger sequencing for *recD2*$^{A227E}$ | CTTGGAATTGCAACTTGTTCATTG |
| Amplify *RS12090* ←/→ *RS12095* for pIMAY* F | TATCGATAAGCTTGATATCGCTATAAAAACTTAGTATTCCAGTTG |
| Amplify *RS12090* ←/→ *RS12095* for pIMAY* R | TGGAGCTCCACCGCGGTGGCTTTAATTTCACTTGTAGGAATTTTG |
| Check pIMAY*-*RS12090* ←/→ *RS12095* integration F | TCTAATGACGCTACCTTCTCTTC |
| Check pIMAY*-*RS12090* ←/→ *RS12095* integration R | TGTATTCGTGGTCTTCAGTATCG |
| Sanger sequencing for *RS12090* ←/→ *RS12095* | TTCCGACCATTAACGCTGTAG |
| Mutagenesis of *lexA* to mutant allele S130A F | TAATATACCAGCCTCAATCATAGCGTCGCCTACGACGTTTAATATG |
| Mutagenesis of *lexA* to mutant allele S130A R | CATATTAAACGTCGTAGGCGACGCTATGATTGAGGCTGGTATATTA |
| Mutagenesis of *lexA* to mutant allele G94E F | TTTCTACTGCGGTAATAGGAACCTCTGCTGTGACTTTACCAATAAC |
| Mutagenesis of *lexA* to mutant allele G94E R | GTTATTGGTAAAGTCACAGCAGAGGTTCCTATTACCGCAGTAGAAA |
| Amplify *lexA* for pIMAY* F | TGGAGCTCCACCGCGGTGGCTTACATTTCGCGGTACAAAC |
| Amplify *lexA* for pIMAY* R | TATCGATAAGCTTGATATCGATGAGAGAATTAACAAAACGAC |
| Sanger sequencing for both *lexA* mutant alleles | TTTCGCGGTACAAACCAATTAC |
| Check pIMAY*-*lexA* integration_F | ATAATGGCTCGCTCCTGTAAAT |
| Check pIMAY*-*lexA* integration_R | TCCCATCTCTTTGTTCGCTATT |
| Check transposon insertion for *recF*::Tn _F | TGAAACACGTCGCGGTAAA |
| Check transposon insertion for *recF*::Tn _R | CATCTCGATGTGGTCCGAATAA |
| Check transposon insertion for *xerC*::Tn _F | TTCTTAAGTCATCAACGAGTAA |
| Check transposon insertion for *xerC*::Tn _R | AAGCGTTACTTGCCCATCTC |
| Check transposon insertion for *recG*::Tn _F | CGCGTTTATGGGCTTGATTG |
| Check transposon insertion for *recG*::Tn _R | GTATCGGCGTTGCTGTCATA |
| Check transposon insertion for *sbcD*::Tn _F | GGAGAATTAGGCGGCATGTT |

| | |
|---|---|
| Check transposon insertion for *sbcD*::Tn _R | ACCTCTCTTTACCATCGTGATTT |
| Check transposon insertion for *recJ*::Tn _F | CCATCCTACTTGTGCACCTAA |
| Check transposon insertion for *recJ*::Tn _R | CTTCAATCTTCATTGCCGTT |
| Check transposon insertion for *recQ*::Tn _F | TCGTAATGTGCTTGGTGTCTTA |
| Check transposon insertion for *recQ*::Tn _R | TGGTAACTCGGCCTGAAATC |
| Check transposon insertion for *recX*::Tn _F | GTTGCGTGTACTTCTTTCGATTT |
| Check transposon insertion for *recX*::Tn _R | TGATCCAGTGCCGAAGATTAC |
| Check transposon insertion for *uvrA*::Tn _F | TGGGTCAGGTAAATCGTCATTAG |
| Check transposon insertion for *uvrA*::Tn _R | AAATCAGAACTCGTCGGTATCC |
| Check transposon insertion for *dinG*::Tn _F | GGCGTCTCTGTAATCCAAATCA |
| Check transposon insertion for *dinG*::Tn _R | GATGCCGCTACTACTGCTAAA |
| Check transposon insertion for *RS09290*::Tn _F | TGTAAGTGAGGAGGGATGTACT |
| Check transposon insertion for *RS09290*::Tn _R | CAATCGCTTGCTTCGTTCTTC |
| Check transposon insertion for *rarA*::Tn _F | ACCAAGAGGCATTATCAGAAGAA |
| Check transposon insertion for *rarA*::Tn _R | AGTTGTAGCACCGATCAAGAC |
| Check transposon insertion for *xseA*::Tn _F | GCACCTGTACTCGCTGTTAAA |
| Check transposon insertion for *xseA*::Tn _R | CTGGCGTTAAAGGTCACGAA |
| Check transposon insertion for *ssbB*::Tn _F | CAAAGTGTGTTTGTCCGTCTTTAT |
| Check transposon insertion for *ssbB*::Tn _R | AATCGTAATTGTCGGGAGACTG |
| Check transposon insertion for *recT*::Tn _F | CTTGCTTGAATGTATCGCCTTTAT |
| Check transposon insertion for *recT*::Tn _R | TTCACCAAGTAATGCCATGAAAC |
| Check transposon insertion for *recQ2*::Tn _F | AAGCACTCGTTGCGACTATAA |
| Check transposon insertion for *recQ2*::Tn _R | AGCGGTTGTCTTAGCATTGA |
| Check transposon insertion for *RS12085*::Tn _F | CCATTACCATTCGGCGTTTC |
| Check transposon insertion for *RS12085*::Tn _R | TGAAGAGAAGGTAGCGTCATTAG |
| Check transposon insertion for *dinB*::Tn _F | CTTCATCATCATTCACGTCTGTTG |
| Check transposon insertion for *dinB*::Tn _R | AGCATCTGCAGGTGTTTCTTA |
| Check transposon insertion for *mfd*::Tn _F | CTGCGTTAGCTCAAGGTAAGA |
| Check transposon insertion for *mfd*::Tn _R | CGCTGCGTTTCAACATCAA |
| Check transposon insertion for *ruvX*::Tn _F | AGCACCTTCTTCAGCTAAGATAAC |
| Check transposon insertion for *ruvX*::Tn _R | GGCACAAGGATTAGACACACTC |
| Check transposon insertion for *RS08985*::Tn _F | AGCGCCAATAACATCTTCTACT |
| Check transposon insertion for *RS08985*::Tn _R | AGTGATCCGACATTGTCCATTTA |
| Check transposon insertion for *RS06660*::Tn _F | TTTGGTGGCGGTGTATTAGG |
| Check transposon insertion for *RS06660*::Tn _R | TGCAGAACGACCAGGTATTG |
| Check transposon insertion for *trhA*::Tn _F | CAACCTATCCAACCACCTACAA |
| Check transposon insertion for *trhA*::Tn _R | AATGCGGCATCTCATGGT |
| Check transposon insertion for *RS12060*::Tn _F | TCCAGGTGGAAGATCGAGTAT |
| Check transposon insertion for *RS12060*::Tn _R | GGTAAAGGTGGTGTCGGTAAA |
| Check transposon insertion for *RS03470*::Tn _F | CCAGCTTGACGACCTTTCATA |
| Check transposon insertion for *RS03470*::Tn _R | GGCGTACGTATACAGCCTTATC |
| Check transposon insertion for *sepA*::Tn _F | ACGTTGTTGCAACTGTGTAAG |
| Check transposon insertion for *sepA*::Tn _R | TTTGTACTTTCTGGTGCGATTT |
| Check transposon insertion for *RS10630*::Tn _F | TCTCTGTCGCTTGTTGTTCTG |
| Check transposon insertion for *RS10630*::Tn _R | AGGGTATGTCGTGTGCAATG |
| Check transposon insertion for *RS03710*::Tn _F | TGCAGAAATGAAACTTGCTGTC |
| Check transposon insertion for *RS03710*::Tn _R | CTACAGCTATAGTGACGATGATT |
| Check transposon insertion for *RS12505*::Tn _F | CTTCTCGTCCAGCATCTGTT |

| | |
|---|---|
| Check transposon insertion for *RS12505*::Tn _R | GTGATTCAATCGCATCGGTTATT |
| Check transposon insertion for *polA*::Tn_F | TCTGGCGTCCACCTTTATATTC |
| Check transposon insertion for *polA*::Tn_R | TGGAGTGCTCATCAAGAACAA |
| Check transposon insertion for *recA*::Tn_F | TCGGTAAAGGTGCCGTAATG |
| Check transposon insertion for *recA*::Tn_F | ACGTAACGCTTGTGACATTAAAC |
| Check transposon insertion for *rexA*::Tn_F | GGTGACACCATACGAAGAAGAA |
| Check transposon insertion for *rexA*::Tn_F | CTCTTGAACTCGGTTCGTATCT |
| Check transposon insertion for *RS12090*::Tn _F | TTCCGACCATTAACGCTGTAG |
| Check transposon insertion for *RS12090*::Tn _R | ACTCTCCATTGAATACGCACTT |
| Check transposon insertion for *RS12095*::Tn _F | TTCCGACCATTAACGCTGTAG |
| Check transposon insertion for *RS12095*::Tn _R | TGTATTCGTGGTCTTCAGTATCG |
| Check transposon insertion for *recD2*::Tn_F | CGGTACCACCCTAGTTATAAATGC |
| Check transposon insertion for *recD2*::Tn_F | ATGCAAGATACAATTTGGGCTTAG |
| Check transposon insertion for *umuC*::Tn_F | GTTGTTGCAGATACTAAGCG |
| Check transposon insertion for *umuC*::Tn_R | AAAGCATTGAACACCATAACCG |
| Gibson assembly for pKK30 F | GCGGCCGCTAGCCTAGGAGC |
| Gibson assembly for pKK30 R | ATCGCCTGTCACTTTGCTTGATATATGA |
| Amplify *xerC* for pKK30 F | CAAGCAAAGTGACAGGCGATAAACTAGAATTAAAGATAAAAAAGAACG |
| Amplify *xerC* for pKK30 R | GCTCCTAGGCTAGCGGCCGCTCATGTTTCATTCTCCTTTTTC |
| Amplify RS12060 for pKK30 F | ATCAAGCAAAGTGACAGGCGATCTATTTATTTGTAGCAGCTATAAC |
| Amplify RS12060 for pKK30 R | GAGCTCCTAGGCTAGCGGCCGCTCTCATTTGCTTTTGTAGC |
| Amplify *recA* for pKK30 F | CAAGCAAAGTGACAGGCGATTTGTATTATCGATAAAAATATAAGCAC |
| Amplify *recA* for pKK30 R | GCTCCTAGGCTAGCGGCCGCCTATTCTTCGTCAAATAATGAC |
| Check junctions in JE2 WT_F | AATTCGCCCACTATAGGATTGG |
| Check junctions in JE2 WT_R1 | GGAGGCCAATGATTGTGAATTA |
| Check junctions in JE2 WT_R2 | TTTGAAACATTGCCGGAAGAAA |
| qPCR for junction in DLX resistant population WT2_F | GCAGTTGCAACACTATCTTTACC |
| qPCR for junction in DLX resistant population WT2_R | CGATGAGTGCTAAGTGTTAGGG |
| qPCR for junction in DLX resistant population WT3_F | TACTCCAGTACTGTATCTTTACAATCG |
| qPCR for junction in DLX resistant population WT3_R | AACTGAATGACAATATGTCAACG |
| qPCR for housekeeping gene *rpoC*_F | CTGTGAAAGAATTTTCGGAC |
| qPCR for housekeeping gene *rpoC*_R | CTTTCACGACGTACTTTAGA |
| qPCR for *sdrM*_F | GCAATGATCGCAATCGGTAT |
| qPCR for *sdrM*_R | GGCATAGTTGGCAGTGTTTG |
| qPCR for *trhA*_F | AGACATACTCACAGACGCAAG |
| qPCR for *trhA*_R | AATGCGGCATCTCATGGT |
| qPCR for *RS12085*_F | TTCTTCTAAGTATCCTGCCTTGTC |
| qPCR for *RS12085*_R | ATTGTGGCGCTTAGTGAAGA |
| qPCR for *RS12090*_F | AGGTTCGATGTCCAAAGATCAA |

| | |
|---|---|
| qPCR for *RS12090*_R | GCAACTCTAGCTAGCTTATTACCC |
| qPCR for *RS12095*_F | GCAGTTCCTAAATTAGACAGCAAAG |
| qPCR for *RS12095*_R | ATGTATTCGTGGTCTTCAGTATCG |

*Underlined sequences represent homology for Gibson assembly.
